# A CSB-PAF1C axis restores processive transcription elongation after DNA damage repair

**DOI:** 10.1101/2020.01.04.894808

**Authors:** Diana van den Heuvel, Cornelia G. Spruijt, Román González-Prieto, Angela Kragten, Michelle T. Paulsen, Di Zhou, Haoyu Wu, Katja Apelt, Yana van der Weegen, Kevin Yang, Madelon Dijk, Lucia Daxinger, Jurgen A. Marteijn, Alfred C.O. Vertegaal, Mats Ljungman, Michiel Vermeulen, Martijn S. Luijsterburg

## Abstract

The coordinated transcription of genes involves the regulated release of RNA polymerase II (RNAPII) from promoter-proximal sites into active elongation. DNA lesions in transcribed strands block elongation and induce a strong transcriptional arrest. The transcription-coupled repair (TCR) pathway efficiently removes transcription-blocking DNA lesions, but this is not sufficient to resume transcription. Through proteomics screens, we find that the TCR-specific CSB protein loads the evolutionary conserved PAF1 complex (PAF1C) onto RNAPII in promoter-proximal regions in response to DNA damage. PAF1C is dispensable for TCR-mediated repair, but is essential for recovery of RNA synthesis after UV irradiation, suggesting an uncoupling between DNA repair and transcription recovery. Moreover, we find that PAF1C promotes RNAPII pause release in promoter-proximal regions and subsequently acts as a processivity factor that stimulates transcription elongation throughout genes. Our findings expose the molecular basis for a non-canonical PAF1C-dependent pathway that restores transcription throughout the human genome after genotoxic stress.

## Introduction

The transcription of protein-coding genes involves RNA polymerase II enzymes (RNAPII), which pull DNA through their active sites and generate nascent transcripts. After initiation at the promoter, the majority of RNAPII molecules in metazoan cells pause at promoter-proximal sites, which is enforced by the negative elongation factors DSIF and NELF (Chen et al., 2018; Vos et al., 2018b). The regulation of RNAPII pause release in response to environmental cues involves positive elongation factors, such as p-TEFb and the PAF1 complex (PAF1C) (Chen et al., 2018; Vos et al., 2018b; Yu et al., 2015). Both PAF1C and DSIF also act beyond pause release by stimulating the acceleration of RNAPII in promoter-proximal regions to ensure processive transcription elongation throughout genes (Fitz et al., 2018; Hou et al., 2019; Kim et al., 2010; Wada et al., 1998).

The presence of bulky DNA damage in the transcribed strand of active genes is a major complication during transcription (Brueckner et al., 2007; Moné et al., 2001). Persistent stalling of RNAPII at DNA lesions is highly toxic and constitutes an efficient trigger for apoptosis (Ljungman and Zhang, 1996). The presence of DNA lesions triggers a genome-wide transcriptional arrest due to stalling of elongating RNAPII at DNA lesions (Brueckner et al., 2007). In addition, UV irradiation also inhibits transcription initiation through the stress-induced transcription repressor ATF3 (Epanchintsev et al., 2017; Kristensen et al., 2013). It is essential that cells overcome this arrest and restore transcription after repair to maintain gene expression.

The transcription-coupled nucleotide excision repair (TCR) pathway efficiently removes transcription-blocking DNA lesions through the ATP-dependent chromatin-remodeling factor CSB / ERCC6 (Citterio et al., 2000) and the CUL4A-based E3 ubiquitin ligase complex containing CSA / ERCC8 (Groisman et al., 2003). Mutations in the *CSB* and *CSA* genes cause Cockayne syndrome, which is characterized by UV sensitivity, premature ageing and severe developmental and neurological dysfunction, possibly caused by the prolonged arrest of RNAPII at DNA lesions (Laugel et al., 2010; Lehmann, 2003) resulting in transcriptional misregulation (Proietti-De-Santis et al., 2006; Wang et al., 2014). CSB is ubiquitylated and degraded in response to DNA damage, which is counteracted by the UVSSA protein (Fei and Chen, 2012; Nakazawa et al., 2012; Schwertman et al., 2012). The stalling of RNAPII and the recruitment of CSB, CSA and UVSSA initiates TCR, which is subsequently followed by the assembly of the TFIIH complex, XPA, and the endonucleases XPG and ERCC1-XPF resulting in removal of the DNA lesion (Marteijn et al., 2014).

Although the TCR-mediated clearing of DNA lesions from the transcribed strands is essential, the precise mechanisms required for recovery of transcription after DNA repair remain unresolved. It has been postulated that stalled RNAPII molecules are reactivated following TCR-mediated repair, which would require re-positioning of the nascent transcript within the active site through hydrolysis to generate a new 3’ end (Donahue et al., 1994). Alternatively, it has been suggested that RNAPII is released from the DNA template (Chiou et al., 2018), followed by transcription recovery from the promoter (Andrade-Lima et al., 2015). Both CSA and CSB are essential for transcription recovery, which could be due to their role in clearing transcription-blocking DNA lesions (Mayne and Lehmann, 1982). In addition, the CS proteins also mediate the proteolytic degradation of ATF3 at later time-points after UV irradiation, thereby eliminating its repressive impact on transcription initiation (Epanchintsev et al., 2017; Kristensen et al., 2013). Furthermore, the histone chaperones HIRA (Adam et al., 2013) and FACT (Dinant et al., 2013) and the histone methyltransferase DOT1L (Oksenych et al., 2013) play important roles in the recovery of transcription. However, the HIRA-dependent deposition of H3.3 and the FACT-mediated exchange of H2A at sites of local UV damage also occur in TCR-deficient cells (Adam et al., 2013; Dinant et al., 2013). Thus, the precise mechanisms involved in transcription recovery and their coordination with TCR-mediated repair remain to be established.

In this study, we define a new transcription recovery pathway that involves the CSB-dependent association of the PAF1 pausing and elongation complex with RNAPII specifically after UV irradiation. We show that PAF1 is not involved in TCR-mediated repair, but specifically regulates RNAPII pause release and elongation activation from promoter-proximal regions. These findings identify an alternative p-TEFb-independent pathway that relies on CSB for the activation of paused RNAPII complexes by PAF1C to restore transcriptional activity and overcome DNA damage-induced silencing throughout the human genome.

## Results

### Identification of the PAF1 complex as a UV-specific interactor of CSB

In order to identify DNA damage-specific interactors of CSB, we stably expressed GFP-tagged CSB in SV40-immortalized CS1AN fibroblasts derived from a Cockayne syndrome B patient. Cells were cultured in light (L) or heavy (H) SILAC medium for 14 days, and subsequently mock-treated (L) or exposed to UV-C light (H). Immunoprecipitation of GFP-CSB from the solubilized chromatin fraction followed by mass spectrometry (MS) identified 172 proteins that showed at least 2-fold stronger association with chromatin-bound CSB in UV-irradiated cells (1 hr) compared with control cells. Among the top interactors were 8 subunits of the RNA polymerase II (RNAPII) complex (Sainsbury et al., 2015), and 4 subunits of the human polymerase-associated factor 1 complex (PAF1C; (Zhu et al., 2005)) (Figure 1A).

**Figure 1.**
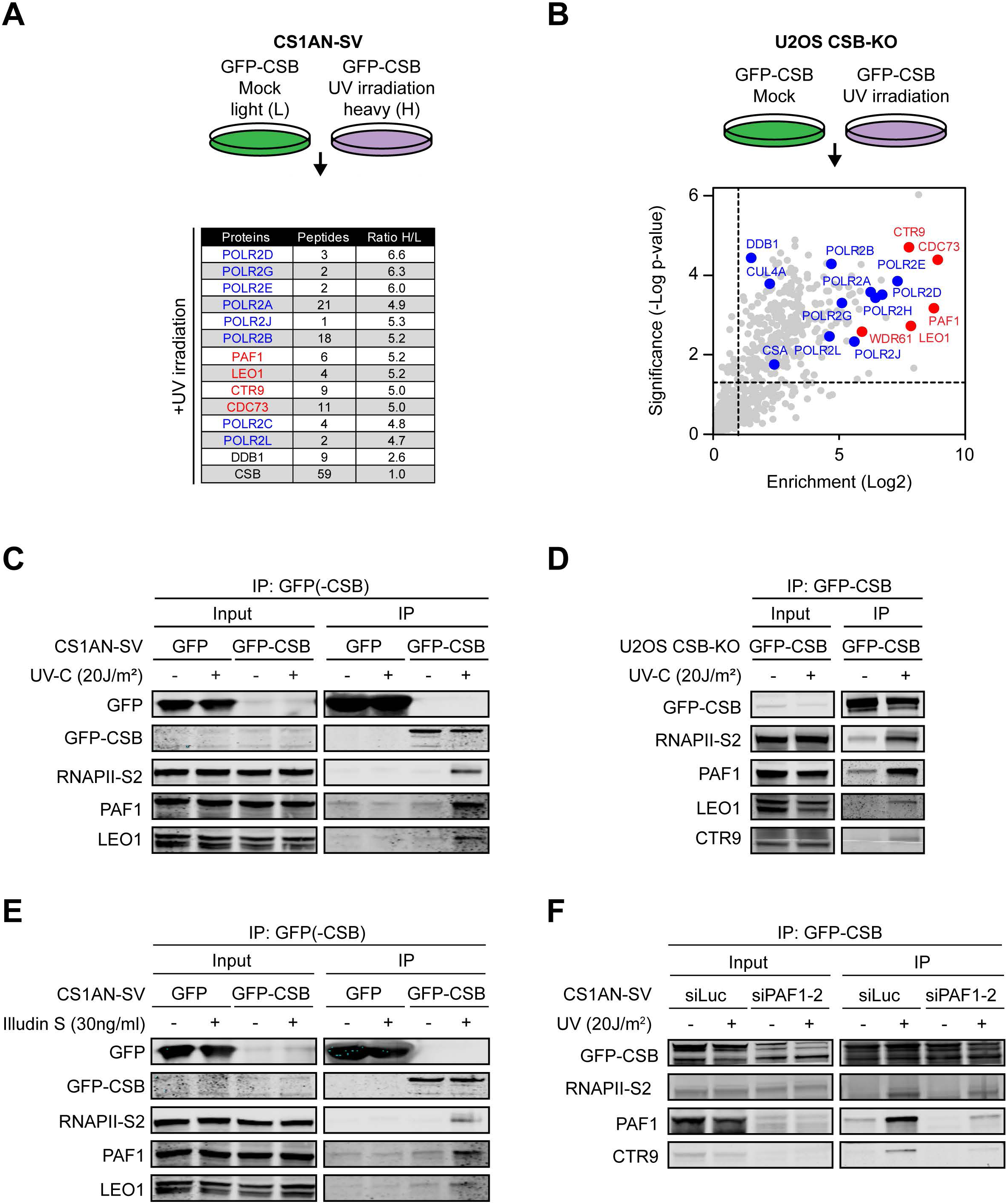
PAF1C is a UV-induced interactor of CSB. (**A**) Results of SILAC-based MS after GFP-CSB pull-down from CS1AN-SV cells. The number of peptides identified and the UV-induced enrichment (ratio H / L) are shown. (**B**) Volcano plot depicting the UV-specific enrichment of proteins after pull-down of GFP-CSB from U2OS CSB-KO cells analysed by label-free MS. The enrichment (log^2^) is plotted on the x-axis and the significance (t-test −log^10^ p-value) is plotted on the y-axis. Highlighted are significantly enriched subunits of RNAPII (blue) and PAF1C or other important proteins (red). (**C**) Co-immunoprecipitation of GFP or GFP-CSB from CS1AN-SV cells in the absence or presence of UV-induced DNA damage. (**D**) Co-immunoprecipitation of GFP-CSB from U2OS CSB-KO cells in the absence or presence of UV-induced DNA damage. (**E**) As in C, but after the induction of DNA damaged by treatment with 30 ng/mL Illudin S for 3hrs. (**F**) As in C, but after transfection with control (siLuc) or PAF1-specific siRNAs.

To confirm these interactions in another cell line, we generated CSB-KO cells in U2OS by CRISPR-Cas9-mediated genome editing, and subsequently re-expressed GFP-CSB in these cells. Expression of GFP-CSB rescued the sensitivity of CSB-KO cells to Illudin S, which causes transcription-blocking DNA lesions, confirming the functionality of the GFP-tagged CSB protein (Supplemental Figure 1A). Label-free quantification proteomics confirmed a strong association of GFP-CSB with RNAPII subunits, the CSA-DDB1-CUL4A complex, and all 5 PAF1C subunits after UV irradiation (Figure 1B). Intensity-based absolute quantification (iBAQ) of protein amounts (Schwanhausser et al., 2011), indicates that at least 60% of the isolated CSB molecules associate with PAF1C and RNAPII subunits after exposure to UV light (Supplemental Figure 1B).

To validate these interactions, we performed co-IP experiments in GFP-CSB-complemented CS1AN-SV cells (Figure 1C) and U2OS CSB-KO cells (Figure 1D), which confirmed that CSB associates with RNAPII as well as with PAF1C subunits PAF1, LEO1 and CTR9 after UV irradiation. Treatment of cells with Illudin S also enhanced the association of CSB with both RNAPII and PAF1C (Figure 1E). To ensure the specificity of the detected bands in our co-IP experiments, we pulled-down GFP-CSB from cells transfected with control or PAF1 siRNAs (Figure 1F). Western blot analysis showed that the knockdown of PAF1 was efficient and also lowered the steady-state levels of CTR9, suggesting destabilization of the PAF1C. Importantly, knockdown of PAF1 severely the UV-induced interaction of CSB with both PAF1 and CTR9 compared to control cells (Figure 1F). In summary, we identify the PAF1 complex as a novel interactor of CSB in response to transcription-blocking DNA damage.

### The PAF1 complex strongly associates with RNAPII and CSB after UV irradiation

PAF1C is an evolutionary conserved pausing and elongation complex that consists of 5 subunits (Figure 2A) (Chen et al., 2015b; Chen et al., 2017; Hou et al., 2019; Yu et al., 2015). Considering that CSB interacts robustly with RNAPII only in the presence of transcription-blocking DNA damage, it is possible that we indirectly detect PAF1C through the UV-induced association of CSB with the RNAPII complex. Alternatively, it is possible that UV-induced DNA damage enhances the association between PAF1C and RNAPII. To distinguish between these possibilities, we generated RPE-hTERT cells stably expressing GFP-tagged LEO1 (Figure 2B). Immunoprecipitation of GFP-LEO1, but not GFP tagged with a nuclear localization sequence (GFP-NLS), showed a robust interaction with PAF1 and CTR9 in both control and UV-exposed cells (Figure 2B). In contrast, we detected CSB and RNAPII in GFP-LEO1 immunoprecipitates only after UV exposure, suggesting that PAF1C readily associates with both CSB and RNAPII in response to UV irradiation (Figure 2B). Increased UV-induced binding of PAF1C with RNAPII was also observed after immunoprecipitation of GFP-tagged CTR9 (Supplemental Figure 1C).

**Figure 2.**
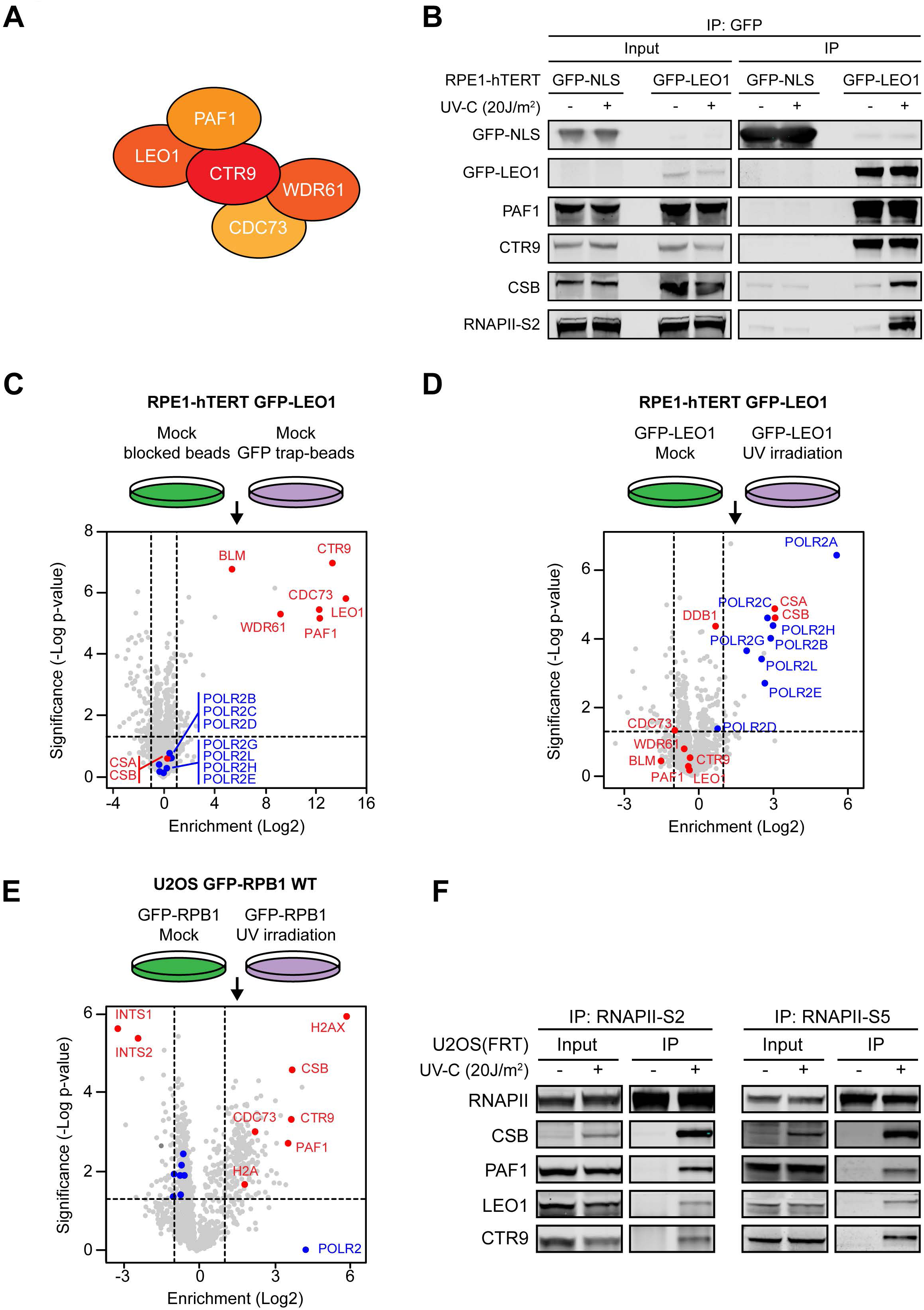
CS proteins and RNAPII show a UV-induced interaction with PAF1C. (**A**) Schematic representation of the PAF1C subunits. (**B**) Co-immunoprecipitation of GFP-NLS or GFP-LEO1 from RPE1-hTERT cells in the absence or presence of UV-induced DNA damage. (**C-E**) Volcano plot depicting the enrichment of proteins after pull-down of (**C**, **D**) GFP-LEO1 from RPE1-hTERT, or (**E**) GFP-RPB1 from U2OS cells. Samples were analysed by label-free MS. The enrichment (log^2^) is plotted on the x-axis and the significance (t-test −log^10^ p-value) is plotted on the y-axis. Highlighted are significantly enriched subunits of RNAPII (blue) and other proteins and the PAF1C (red). (**F**) Co-immunoprecipitation of endogenous RNAPII using Ser2- or Ser5-specific antibodies from U2OS cells in the absence or presence of UV-induced DNA damage.

To confirm these results in an unbiased manner, we performed label-free mass spectrometry on the RPE1 cells expressing GFP-LEO1. Following GFP-LEO1 pull-down from unirradiated cells, we detected strong interactions with all PAF1C subunits (PAF1, CTR9, WDR61, and CDC73), while the RNAPII complex or CS proteins were not significantly enriched (Figure 2C). Exposure of cells to UV light did not affect the interactions between PAF1C subunits (Supplemental Figure 1D), but did trigger the association with at least 8 subunits of the RNAPII complex as well as with CSA and CSB (Figure 2D). Based on the iBAQ values, we estimate that between 1-3% of the isolated LEO1 proteins associates with CS proteins and RNAPII subunits in response to UV irradiation, while this was ∼0.2% in unirradiated cells (Supplemental Figure 1E).

To further study these interactions, we performed targeted immunoprecipitation and unbiased label-free proteomics on U2OS cells stably expressing GFP-tagged RPB1 (Caron et al., 2019), which is the largest subunit of the RNAPII complex. Label-free quantification proteomics in unirradiated cells revealed interactions with all 11 other subunits of the RNAPII complex and many known interactors of RNAPII, including 22 Mediator subunits (Soutourina, 2018), RECQL5 (Saponaro et al., 2014), and WWP2 (Caron et al., 2019) (Supplemental Figure 1F). In line with our previous findings, immunoprecipitation of GFP-RPB1 in unirradiated cells revealed no or only a very weak interaction with PAF1C subunits and no interaction with CSB (Supplemental Figure 1G, H). Conversely, immunoprecipitation of GFP-RPB1 from UV-irradiated cells revealed a strong association of RNAPII with PAF1, LEO1, CTR9 and CSB by label-free quantification proteomics (Figure 2E) and Western blot analysis (Supplemental Figure 1H). Based on the iBAQ values, we estimate that UV-induced DNA damage triggers a ∼70-fold increase in the association between PAF1C and RNAPII, which is roughly comparable with the UV-C-induced increase we detect between RNAPII and CSB (Supplemental Figure 2A, B). Immunoprecipitating of endogenous RNAPII (RNAPIIo) using either a Ser5-P or a Ser2-P-specific antibody confirmed that UV irradiation triggered a strong association of PAF1C and CSB with RNAPIIo, while these interactions were largely absent in non-irradiated cells (Figure 2F).

The association of PAF1 with RNAPII during transcription elongation requires the activity of CDK9 in the pTEFb kinase complex (Vos et al., 2018a; Yu et al., 2015). However, inhibition of CDK9 did not affect the UV-induced association of PAF1 with RNAPII (Supplemental Figure 2C), suggesting that this UV-induced interaction is established differently from the canonical interaction during transcription elongation. In summary, our findings reveal a strong UV-induced association of PAF1C with RNAPII and CSB.

### The UV-induced interaction between PAF1C and RNAPII is mediated by CSB

We next asked whether the UV-induced association of PAF1C with RNAPII was dependent on CSB. To address this, we generated wild-type or CSB-KO cells stably expressing GFP-PAF1. Immunoprecipitation of GFP-PAF1 showed a constitutive association with CTR9, while UV irradiation triggered a strong association with CSB, CSA, and RNAPIIo. Strikingly, knockout of CSB did not affect the association of PAF1 with CTR9, but it prevented the UV-induced association of PAF1 with both CSA and RNAPIIo (Figure 3A).

**Figure 3.**
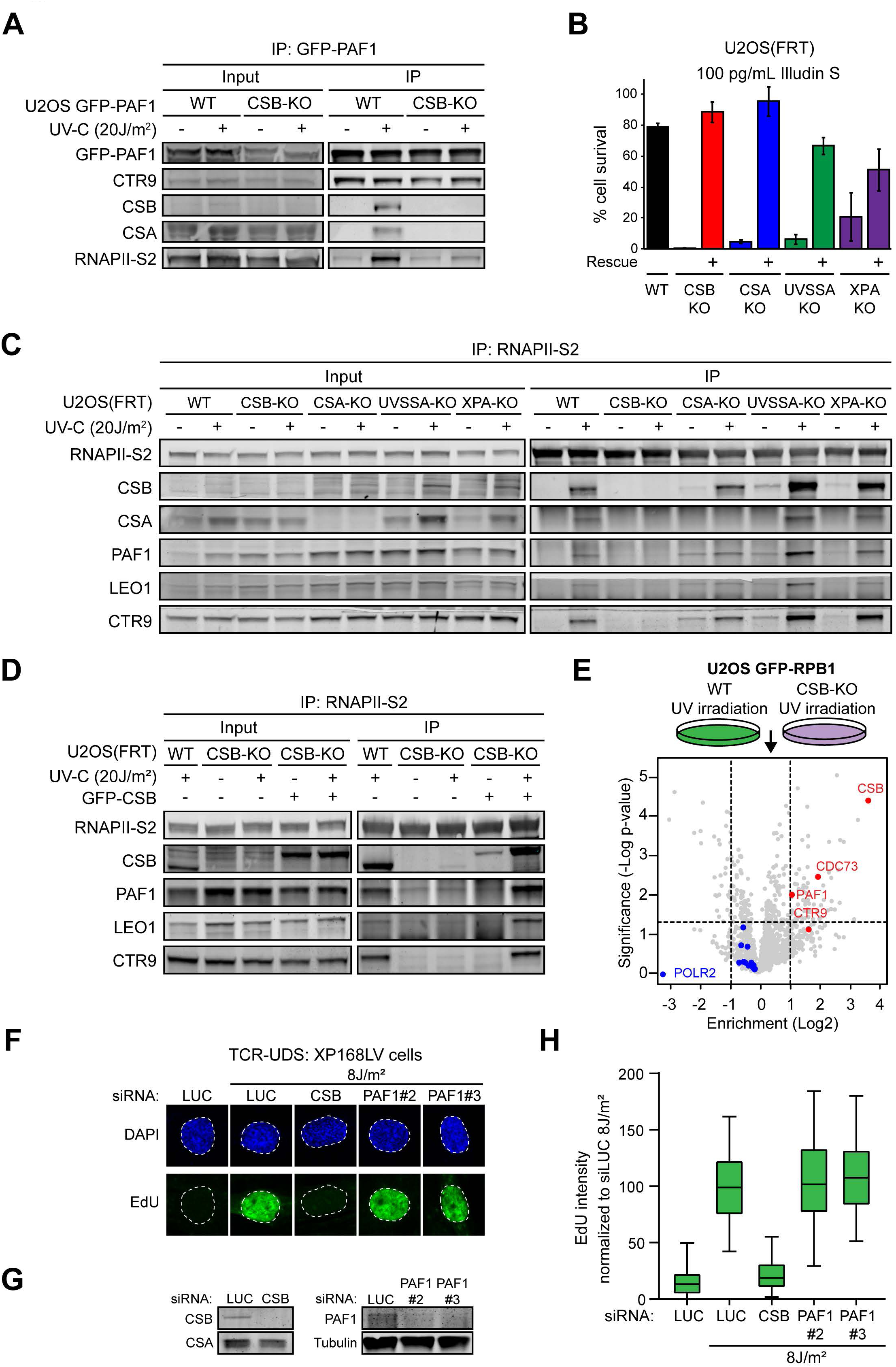
The RNAPII – PAF1C interaction is mediated by CSB and not involved in repair. (**A**) Co-immunoprecipitation of GFP-PAF1 from U2OS WT and U2OS CSB-KO cells in the absence or presence of UV-induced DNA damage. (**B**) Clonogenic Illudin S survival of WT, CSA, CSB, and UVSSA knockout and rescue cell lines at a single dose. Data represent mean ± SEM of at least two independent experiments. (**C**) Co-immunoprecipitation of endogenous RNAPII-S2 from U2OS cells (WT or indicated KO cells) in the absence or presence of UV-induced DNA damage. (**D**) As in C, but now also included U2OS CSB-KO cells reconstituted with GFP-CSB. (**E**) Volcano plot depicting the UV-induced and CSB-specific enrichment of proteins after pull-down of GFP-RPB1 from U2OS WT versus U2OS CSB-KO cells analysed by label-free MS. The enrichment (log^2^) is plotted on the x-axis and the significance (t-test −log^10^ p-value) is plotted on the y-axis. Highlighted are significantly enriched subunits of PAF1C and CSB itself (red), or RNAPII (blue). (**F**) TCR-UDS assay in XP-C primary fibroblasts (XP168LV) during 8 hrs following UV in cells transfected with the indicated siRNAs. (**G**) Validation of the knockdown in F by Western blot analysis. (**H**) Quantification of TCR-UDS signal from F. Data is represented in boxplots showing the median, 50% and 95% percentile of at least two independent experiments.

The assembly of TCR complexes fully depends on CSB. It is therefore possible that another TCR protein that acts downstream of CSB mediates the UV-induced interaction between PAF1C and RNAPII. To investigate this possibility, we generated a collection of TCR knock-outs using CRISPR-Cas9-mediated genome editing in both parental and GFP-RPB1-expressing U2OS cells.

Knock-out of UVSSA was confirmed by DNA sequencing due to a lack of specific antibodies (Supplemental Figure 2D), while knock-out of CSB, CSA (Figure 3A-C) and XPA (Supplemental Figure 2E) was confirmed by western blot analysis. Knock-out clones were highly sensitive to Illudin S (Figure 3B; Supplemental Figure 2F), in line with a specific defect in TCR, which was fully rescued by stable expression of GFP-CSB, CSA-GFP, UVSSA-GFP or GFP-XPA in the corresponding knock-out lines (Figure 3B). Immunoprecipitation of endogenous RNAPIIo using Ser2-P antibodies revealed a robust UV-specific interaction with PAF1C in cells deficient for CSA, UVSSA and XPA, while this interaction was abolished in CSB-KO cells (Figure 3C). A similar result was obtained using antibodies against Ser5-P-modified RNAPII (Supplemental Figure 2G). Immunoprecipitation of GFP-RPB1 from the generated knock-out cells confirmed that the UV-induced association of PAF1C with RNAPII was unaffected in CSA-KO and UVSSA-KO cells, but was completely lost in CSB-KO cells (Supplemental Figure 2H). Importantly, re-expression of GFP-tagged CSB in CSB-KO cells at physiological levels restored the association between RNAPII and PAF1C after UV irradiation (Figure 3D).

To further confirm these findings, we performed label-free proteomics on immunoprecipitated GFP-RPB1 from either WT or CSB-KO cells to identify CSB-dependent interactors of RNAPII after UV in an unbiased manner. This approach confirmed that PAF1C subunits CDC73, CTR9 and PAF1 were among the differential interactors between WT and CSB-KO cells (Figure 3E). These findings demonstrate that CSB is specifically required to establish the UV-induced association between PAF1C and RNAPII.

### The PAF1 complex is not required for clearing DNA lesions by TCR

CSB strongly associates with RNAPIIo following UV irradiation and initiates TCR-mediated repair of DNA lesions on actively transcribed strands. Considering the strong CSB-dependent association of PAF1C with RNAPIIo, we asked whether PAF1C has a direct role in TCR. To address this possibility, we monitored unscheduled DNA synthesis (UDS) after global UV-C irradiation as a measure for DNA repair. To specifically capture CSB-dependent repair by TCR, we employed non-dividing primary XP-C patient-derived fibroblasts, which are deficient in global genome repair (GGR; Figure 3F-H). These XP-C cells were globally irradiated with UV-C light (8 J/m^2^) and immediately pulse-labelled for 8 hours with the nucleotide analogue 5-ethynyl-deoxy-uridine (EdU). TCR-specific UDS was visualized using Click-It chemistry combined with tyramide-based signal amplification (Wienholz et al., 2017). Robust incorporation of EdU could indeed be detected in UV-irradiated XP-C cells, which was virtually absent from non-irradiated controls cells (Figure 3F-H). Knockdown of CSB with specific siRNAs prevented the incorporation of EdU demonstrating that our experimental approach reflects TCR-specific repair synthesis. Importantly, knockdown of PAF1 with two independent siRNAs, which was validated by Western blot analysis (Figure 3G), did not affect TCR-specific repair synthesis (Figure 3F-H). These findings demonstrate that PAF1 is not involved in clearing DNA lesions by TCR.

### The PAF1 complex specifically promotes transcription recovery after UV irradiation

The presence of UV-induced DNA lesions triggers the stalling of RNAPII leading to a strong transcription arrest. Several hours following UV irradiation, when transcription-blocking lesions have been removed, TCR-proficient cells start to resume transcription, which does not occur in TCR-deficient cells (Andrade-Lima et al., 2015; Mayne and Lehmann, 1982). In light of the known role of PAF1C in transcription, we sought to address whether PAF1C plays a role in transcription recovery after repair. To this end, we visualized nascent transcription by 5-ethynyl-uridine labeling following global UV irradiation of U2OS cells to measure RNA recovery synthesis (RRS; Figure 4A). Nascent transcription was strongly inhibited at 3 hours after UV irradiation in all conditions (Figure 4B-E). However, transcription significantly recovered 18 hours after UV irradiation in controls cells, but not in TCR-deficient XPA knockdown cells (Figure 4B-E). In order to monitor the role of PAF1 in this process, we used two independent siRNAs, which efficiently lowered PAF1 steady-state protein levels (Figure 4C). Interestingly, PAF1 knockdown significantly impaired the ability of cells to overcome the transcriptional arrest following UV irradiation (Figure 4B, E). To ensure that this phenotype is not due to an off-target effect, we stably expressed an inducible siRNA-resistant version of GFP-tagged PAF1 (Figure 4D), which fully restored RNA synthesis after UV in cells depleted for endogenous PAF1 (Figure 4B, E).

**Figure 4.**
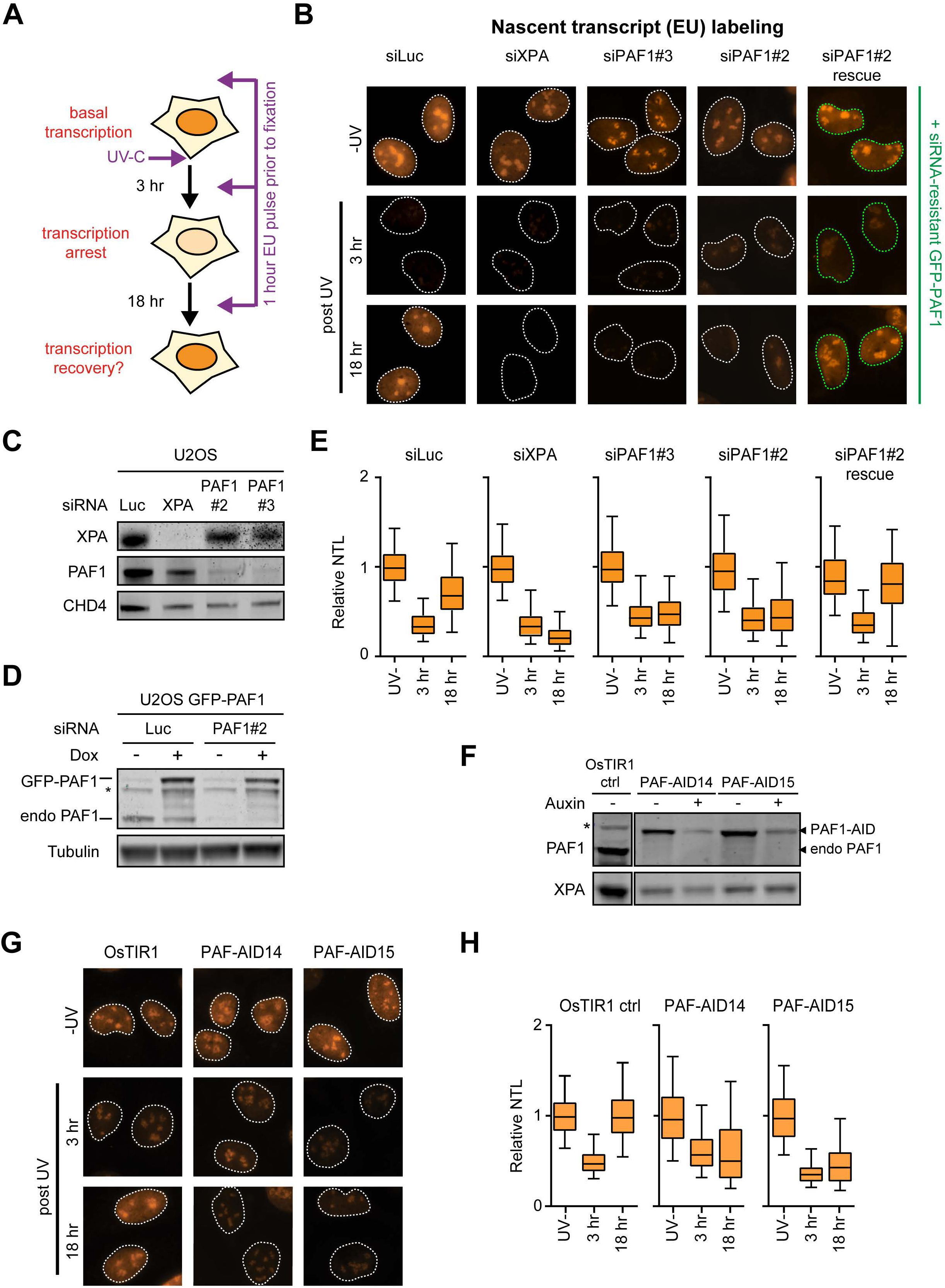
PAF1C loss impairs transcription recovery after UV irradiation. (**A**) Experimental outline of the RRS approach. (**B**) Representative images of U2OS cells transfected with the indicated siRNAs after pulse-labelling with 5-ethynyl-uridine (5-EU) and detection by Click-It chemistry at different time-points after UV. Cells with green outlines express GFP-tagged siRNA-resistant PAF1. (**C**) Validation of the knockdown in B by Western blot analysis. (**D**) Validation of the replacement strategy in B with knockdown of endogenous and simultaneous ectopic expression of siRNA-resistant PAF1 by Western blot analysis. (**E**) Quantification of RRS signal from B. Data is represented in boxplots showing the median, 50% and 95% percentile of three independent experiments. (**F**) Validation of auxin-induced degradation of endogenous PAF1 in two independent U2OS PAF1-AID knockin clones. (**G**) Representative images of U2OS osTIR1 control or two PAF1-AID clones treated with auxin after pulse-labelling with 5-ethynyl-uridine (5-EU) and detection by Click-It chemistry at different time-points after UV. (**H**) Quantification of RRS signal from G. Data is represented in boxplots showing the median, 50% and 95% percentile of four independent experiments.

We next set out to validate these findings in an siRNA-independent manner. Although we initially attempted to generate PAF1 knock-out cells using CRISPR-Cas9-mediated genome editing, this approach was unsuccessful likely due to the fact that PAF1C genes are essential. To circumvent this issue, we first knocked in the rice-specific F-box gene *osTIR1* in the *AAVS1* locus (Natsume et al., 2016). Subsequently, we knocked in an auxin-inducible degron (AID) into both alleles of the endogenous *PAF1* locus to enable conditional protein depletion (Supplemental Figure 3A). Single knockin clones showed strong auxin-induced depletion of PAF1 within 5 hours (Figure 4F). Quantification of nascent transcription in two independent PAF1-AID clones showed that these cells failed to restore transcription after UV irradiation, while osTIR1 control cells treated with auxin showed full transcription recovery (Figure 4G-H). Altogether, these findings reveal an important role of PAF1C in regulating recovery of transcription following genotoxic insult.

### The interaction between PAF1 and RNAPII after UV takes place in promoter-proximal regions

Our finding that PAF1 is not involved in TCR, but is essential for transcription recovery raises the question where the interaction between PAF1, CSB, and RNAPII takes place in the genome. To gain further insight into this, we tested how various transcription inhibitors affected these UV-induced interactions in GFP-RPB1-expressing cells. Treatment of cells with the selective CDK9 inhibitor LDC00067 between 3 and 8 hours (Albert et al., 2014; Garriga and Grana, 2014), which inhibits RNAPII pause release, but not transcription initiation, did not affect the interaction between RNAPII and PAF1C (Figure 5A), which was detected within the first 8 hrs after UV (Figure 5B). Conversely, inhibition of transcription by the more general CDK inhibitor flavopiridol, which inhibits both pause release (through CDK9) and transcription initiation (through CDK7) (Albert et al., 2014; Sedlacek et al., 1996), reduced the binding of PAF1C to RNAPII (Figure 5A). This effect was phenocopied by inhibiting transcription initiation using the selective TFIIH-inhibitor triptolide in a time-dependent manner (Chen et al., 2015a) (Figure 5A; Supplemental Figure 3B). Importantly, control experiments confirmed that all these transcription inhibitors severely reduced the incorporation of 5-EU (Supplemental Figure 3C). These findings suggest that the UV-induced interaction between PAF1C and RNAPII takes place in promoter-proximal regions.

**Figure 5.**
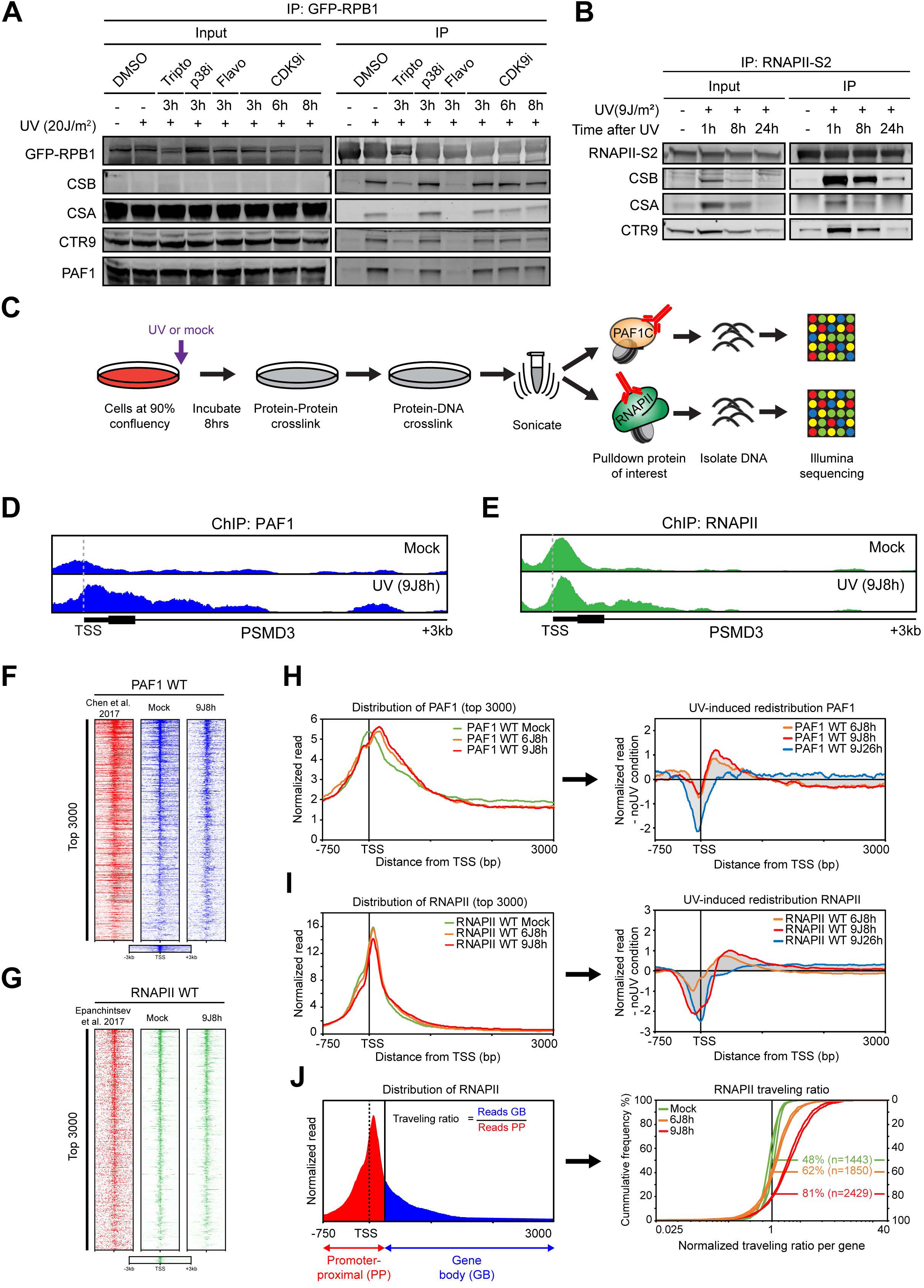
ChIP-seq reveals UV-induced repositioning of PAF1 and RNAPII into promoter-proximal regions. (**A**) Co-immunoprecipitation of GFP-RPB1 in the presence of the indicated inhibitors: 0.55 µM Triptolide (TFIIH/XPB inhibitor), 10 µM SB203580 (p38i), 10 µM Flavopiriol (CDK9 and CDK7 inhibitor), and 10 µM LDC00067 (specific CDK9 inhibitor) for the indicated time. All transcription inhibitors reduced nascent transcription measured by 5-EU incorporation (Supplemental Figure 3C). (**B**) Co-immunoprecipitation of endogenous RNAPII-S2 from U2OS cells at different time-points after UV. (**C**) Outline of the ChIP-seq approach to map PAF1- and RNAPII-binding sites. (**D, E**) UCSC genome browser track showing the read density of (**D**) PAF1 and (**E**) RNAPII signal across the *PSMD3* gene in unirradiated and UV-irradiated cells. (**F**) Heatmaps from PAF1 ChIP-seq data around the transcription start sites (TSS) of the top 3,000 genes that bind PAF1 (identified in Supplemental Figure 3E). Data is ranked based on the PAF1 signal in unirradiated cells (mock; in blue), and compared to published PAF1 ChIP-seq data (in red), and PAF1 ChIP-seq at 8 hrs after 9J/m^2^ UV irradiation (in blue; 9Jh8). (**G**) Heatmaps from RNAPII ChIP-seq data around the TSS of genes as in F. Data of unirradiated cells (mock; in green) are compared to published RNAPII ChIP-seq data (in red), and RNAPII ChIP-seq at 8 hrs after 9J/m^2^ UV irradiation (in green; 9Jh8). (**H**) Metaplots of PAF1 ChIP-seq of the top 3,000 genes around the TSS in unirradiated (mock) and UV-irradiated cells (8h after 6 and 9 J/m^2^). The right panel shows the UV-induced redistribution of PAF1 calculated by subtracting the mock from the +UV distribution profiles for 6 J/m^2^ at 8 hrs, and for 9 J/m^2^ at 8 and 26 hrs. (**I**) As in H, but for RNAPII. (**J**) Schematic representation of the traveling ratio of RNAPII, which is calculated by dividing the reads of the gene body (+250bp to +3kb; blue) over the reads in the promoter-proximal region (- 750bp to +250bp; red). The right panel shows the ratio of the RNAPII traveling ratio (or the normalized traveling ratio) for 3,000 genes relative to the average traveling ratio in the unirradiated control (set to 1). Shown are 3 independent replicates in unirradiated cells (green), and two replicates after UV irradiation with 6 J/m^2^ (in orange) and two replicates after 9 J/m^2^ (in red). The y axes indicate percent of all genes. Percentages and n indicated in the plot refer to the percentage and number of the 3,000 genes with a normalized traveling ratio above 1.

### Genome-wide repositioning of PAF1 and RNAPII at the TSS in response to UV irradiation

One intriguing possibility is that transcription recovery after repair involves the activation of paused RNAPII from promoter-proximal regions. In particular because PAF1C regulates RNAPII pause release during transcription and subsequently stimulates processive elongation throughout genes (Chen et al., 2015b; Chen et al., 2017; Hou et al., 2019; Vos et al., 2018a; Yu et al., 2015).

To gain more insight into where PAF1C binds in response to UV irradiation, we mapped PAF1 chromatin-binding sites in the genome using ChIP-sequencing (ChIP-seq). Double crosslinking using formaldehyde and the protein-protein crosslinker disuccinimidyl glutarate (DSG) was used to preserve chromatin-protein interactions as well as protein-protein interactions within chromatin-bound multi-protein complexes (Figure 5C) (Spruijt et al., 2016). Mapping of sequence reads to the human genome (Hg38) revealed that PAF1 was bound predominantly to transcription start sites (TSSs) and downstream from transcription termination sites (TTSs) (Figure 5D-E; Supplemental Figure 3D). Heatmaps of the distribution of reads around TSSs revealed a large degree of overlap between PAF1 chromatin-binding sites mapped by us compared to published ChIP-seq data for PAF1 that we analyzed in parallel (Chen et al., 2017). From the set of 8,811 genes, we could clearly detect robust binding of PAF1 to the TSS in a subset of ∼3,000 genes (Figure 5F; Supplemental Figure 3E), which also displayed PAF1 binding downstream from the TTSs (Supplemental Figure 3F).

We next sought to address whether the genome-wide binding of PAF1 is altered following UV irradiation. To this end, we globally irradiated cells with a non-lethal dose of UV-C (6 J/m^2^ or 9 J/m^2^), which leads to maximal transcription inhibition at 3 hrs after UV irradiation, and near full transcription recovery in TCR-proficient cells at 18-24 hours after UV irradiation (Fig 4E, H). We decided to perform ChIP-seq experiments at 8 hours after UV since cells already start to resume transcription, but have not yet fully completed transcription recovery at this time point (Andrade-Lima et al., 2015). Immunoprecipitation of RNAPIIo indeed revealed a strong interaction with PAF1C subunit CTR9 at 8 hours after UV, which was lost at 24 hours after UV when cells have fully recovered transcription (Figure 5B). For the ChIP-seq analysis, we focused on a subset of non-overlapping genes and we excluded short genes (< 3kb) that might not be damaged under our experimental conditions (Perdiz et al., 2000), or long genes (> 100kb) that are likely to contain multiple DNA lesions and may show slower recovery. Interestingly, PAF1 became more restricted to the TSS region after UV irradiation (both 6 J/m^2^ and 9 J/m^2^), and showed substantially reduced binding in gene bodies and downstream of the TTS (Supplemental Figure 3G). However, we noted a marked shift in PAF1 binding away from the promoter in the first ∼1 kb downstream of the TSS at 8 hrs after UV, which was lost at 26 hrs after UV (Figure 5H). This region correspond to the promoter-proximal pause site where RNAPII is released under the tight control of regulatory factors in response to environmental cues (Chen et al., 2018). Interestingly, PAF1C is an important regulator of RNAPII pause release (Chen et al., 2015b; Chen et al., 2017; Vos et al., 2018a; Yu et al., 2015), suggesting that PAF1C may release RNAPII from pause sites as a mechanism to restart transcription after UV irradiation.

### A genome-wide shift of RNAPII into gene bodies after UV irradiation

To explore the possibility that PAF1C regulates pause release near TSS sites after UV irradiation, we mapped RNAPII-binding sites by ChIP-sequencing. Heatmaps of the RNAPII distribution in unirradiated cells showed a high degree of overlap with published RNAPII ChIP-seq data (Figure 5G, Supplemental Figure 3H) (Epanchintsev et al., 2017). Importantly, RNAPII most strongly associated with the TSS as well as downstream of the TTS in the same subset of 3,000 genes that were also bound by PAF1 (Figure 5G and Supplemental Figure 3H-I).

Strikingly, UV irradiation triggered a dose-dependent release of RNAPII into the first ∼2 kb downstream of the TSS at 8 hrs after UV irradiation (Figure 5I), which coincided with the region to which a UV-induced shift in PAF1 binding was also observed (Figure 5H). Importantly, the shift in RNAPII binding was no longer detected at 26 hrs after UV irradiation (Figure 5I), which coincides with the loss of PAF1C binding to RNAPII (Figure 5B), and the near complete recovery of nascent transcription in RRS experiments around this time after UV irradiation in control cells (Figure 4E, H). Representative examples of short (ARF6, 5 kb), intermediate (NDUFS5, 10kb) and longer genes (PSMD3, 24 kb) all show the UV-induced redistribution of both PAF1 and RNAPII within the first 2 kb of the gene (Supplemental Figure 4A-C). Importantly, we observe that both RNAPII and PAF1 show a very comparable shift, suggesting that this may reflect the UV-induced loading of PAF1 onto RNAPII in promoter-proximal regions.

To further quantify the release of RNAPII after UV irradiation, we calculated the traveling ratio for each gene, which was defined as the density of RNAPII reads within the first 3kb of the gene body relative to the density in the promoter-proximal region (Figure 5J). An increased traveling ratio indicates that more RNAPII molecules have shifted into the gene body. To allow a direct comparison between our different conditions, we normalized the traveling ratio for each gene after UV irradiation to the traveling ratio of the same gene in unirradiated cells to provide a normalized RNAPII traveling ratio for individual genes. This quantification revealed a dose-dependent increase of the normalized traveling ratio after UV irradiation (Figure 5J), indicating that the shift of RNAPII into the first 2kb of the gene body is a genome-wide phenomenon and not the result from single genes affecting overall binding profiles. Importantly, independent repeats of these different ChIP-seq conditions showed a highly similar shift in the normalized traveling ratios (Figure 5J, Supplemental Figure 4D), suggesting that these changes in RNAPII binding across the genome reflect a very robust cellular response.

### Genes with repositioned PAF1 and RNAPII at the TSS are bound by CS proteins and ATF3

Given that the interaction between PAF1 and RNAPII after UV irradiation is fully dependent on CSB (Figure 3), we asked how the genome-wide binding profiles of PAF1 and RNAPII correlate with those of CSB and CSA. To this end, we re-analyzed genome-wide chromatin-binding profiles of the CS proteins based on recently published data (Epanchintsev et al., 2017). This analysis revealed that CSB is already bound at the TSS of the top 3,000 genes that also have both PAF1 and RNAPII bound in unirradiated cells, while UV irradiation had no appreciable impact on CSB binding to the TSS of these genes (Figure 6A; Supplemental Figure 5A). While initially not bound to these genes in unirradiated cells, CSA became strongly enriched in the top 3,000 PAF1 genes at 8 hrs after UV irradiation (Figure 6A; Supplemental Figure 5A). The striking co-localization of CS proteins and PAF1 chromatin-binding sites in the genome after UV irradiation (Figure 6A) is in line with our interaction data showing that both CS proteins show a strong UV-induced association with the PAF1 complex (Figure 2, Figure 3). Interestingly, after UV irradiation, the top 3,000 genes from our analysis became strongly bound by ATF3 (Figure 6A; Supplemental Figure 5A), which is a repressor that inhibits transcription initiation after UV irradiation (Epanchintsev et al., 2017). These findings suggest that the top 3,000 genes from our analysis are subjected to ATF3-mediated transcriptional repression after UV irradiation, providing a rationale for why these genes require reactivation of transcription by PAF1 from promoter-proximal sites.

**Figure 6.**
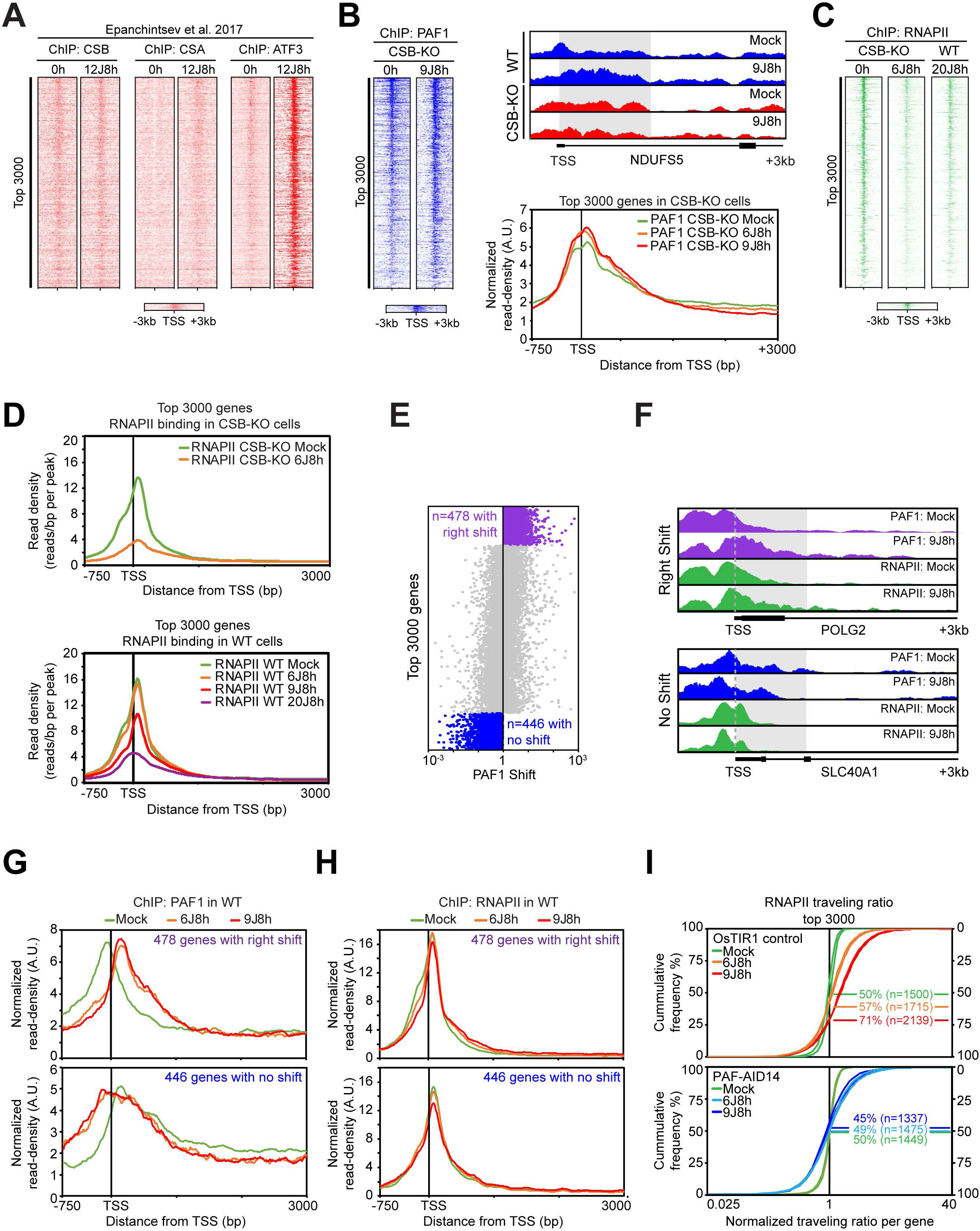
The UV-induced repositioning of RNAPII into promoter-proximal regions requires PAF1C. (**A**) Heatmaps around the TSS from ChIP-seq data on CSB, CSA and ATF3 at 0 or 8 hrs after 12 J/m^2^ UV. The data is re-analysed from the Egly lab and shown for the top 3,000 genes that bind PAF1 (identified in Figure 5F). (**B**) Heatmaps around TSS from PAF1 ChIP-seq data of the top 3,000 genes that bind PAF1 in CSB-KO cells at 0 and 8 hrs after 6 or 9J/m^2^ UV. The right upper panel shows a UCSC genome browser track showing the read density of PAF1 signal across the *NDUFS5* gene in unirradiated and UV-irradiated WT (blue) and CSB-KO cells (red). The right lower panel shows metaplots of PAF1 ChIP-seq of the top 3,000 genes in unirradiated or UV-irradiated (6 and 9 J/m^2^) CSB-KO cells. (**C**) Heatmaps around the TSS from ChIP-seq data on RNAPII of the top 3,000 genes that bind PAF1. Heatmaps are show for CSB-KO cells (at 0 and 8 hrs after 6 J/m^2^) and UV-irradiated WT cells at 8 hrs after 20 J/m^2^. (**D**) Upper panel shows non-normalized metaplots around the TSS of RNAPII ChIP-seq of the top 3,000 genes in unirradiated (mock) or UV-irradiated (8h after 6 J/m^2^) CSB-KO cells showing differences in total RNAPII binding in different conditions. The lower panel shows non-normalized metaplots of RNAPII ChIP-seq of the top 3,000 genes in unirradiated (mock) or UV-irradiated (8h after 6, 9, and 20 J/m^2^) WT cells. (**E**) Quantification of the traveling ratio (or shift) in PAF1 binding after UV. All replicates and UV doses (6 and 9 J/m^2^) were pooled for this analysis, which revealed a set of 478 genes that shows a strong UV-induced shift into promoter-proximal regions in all replicate experiments at 8h after 6 and 9 J/m^2^ (right-shift) and a set of 446 genes that do not show a UV-induced shift in any of the replicate experiments at 8h after 6 and 9 J/m^2^ (no-shift). (**F**) UCSC genome browser track showing the read density of PAF1 and RNAPII signal across the *POLG2* gene (right-shift) and *SLC40A1* gene (no-shift) in unirradiated (mock) and UV-irradiated cells. (**G**) Metaplots of PAF1 ChIP-seq around the TSS of the right-shift 478 genes (upper panel) or the no-shift 446 genes (lower panel) in unirradiated (mock) and UV-irradiated cells (8h after 6 and 9 J/m^2^). (**H**) As in D, but for RNAPII. (**I**) The ratio of the RNAPII traveling ratio (or the normalized traveling ratio) for 3,000 genes relative to the average traveling ratio in the unirradiated control (set to 1) shown for osTIR1 (upper panel) or PAF1-AID clone 14 (lower panel). Shown are 2 independent replicates in unirradiated cells, and two replicates after UV irradiation with 6 J/m^2^, and two replicates after 9 J/m^2^. The y-axes indicate percent of all genes. Percentages and n indicated in the plot refer to the percentage and number of the 3,000 genes with a normalized traveling ratio above 1.

### The repositioning of PAF1 after UV irradiation is stimulated by CSB

Given that PAF1 and CSB strongly interact in response to UV irradiation, we asked whether the repositioning of PAF1 away from TSS sites is dependent on CSB. To address this, we mapped chromatin-binding sites of both PAF1 and RNAPII by ChIP-sequencing in CSB knockout (KO) cells. In unirradiated cells, we did not detect major differences in PAF1 binding to promoter regions in the genomes of WT or CSB-KO cells (compare Figure 6B to Figure 5F). Importantly, UV irradiation did not result in an obvious repositioning of PAF1 downstream of the TSS in CSB-KO cells (Figure 6B), in contrast to our data in WT cells (Figure 5H). These data suggest that the UV-induced association of CSB with PAF1 may be required to drive the genome-wide repositioning of PAF1 in response to UV irradiation.

While loss of CSB did not appreciably affect the binding of RNAPII to TSS sites compared to WT cells without irradiation (Figure 6C, D compare to Figure 5G, I), we detected strongly reduced binding of RNAPII at 8 hrs after UV irradiation with 6J/m^2^ in CSB-KO cells (Figure 6C, D), which was not observed under similar conditions in WT cells (Figure 6D). Strikingly, absolute RNAPII read densities at TSS sites after UV irradiation with 6J/m^2^ in CSB-KO cells were comparable with RNAPII read densities in WT cells after irradiation with a lethal dose of 20J/m^2^ (Figure 6D). These findings suggest that CSB-KO cells through a combination of their inability to repair transcription-blocking lesions and remove ATF3 from TSS sites (Epanchintsev et al., 2017), show a very strong transcriptional repression at 8 hrs after UV irradiation. Of interest, the reduced binding of RNAPII in CSB-KO cells after UV irradiation did not coincide with reduced binding of PAF1 to the same genes (Figure 6B-D), suggesting that these events are unrelated. Furthermore, our data suggest that the repositioning of PAF1 after UV irradiation in the genome is stimulated by CSB in line with the strong UV-induced association between these proteins in our interaction studies (Figures 1-3).

### The release of RNAPII after UV irradiation is dependent on PAF1

To study if the observed release of RNAPII after UV irradiation (Figure 5I) is triggered by PAF1, we first analyzed the correlation between the UV-induced repositioning of PAF1 and RNAPII across genes. To this end, we quantified the relative distribution of PAF1 binding in individual genes by calculating the ratio of reads in the first 1 kb downstream of the TSS relative to the reads immediately upstream or at the TSS (Figure 6E). The distribution profile of PAF1 at 8 hrs after UV irradiation with either 6J/m^2^ or 9J/m^2^ was normalized to the distribution in unirradiated cells to provide a quantification of the relative redistribution of PAF1 in individual genes (Figure 6E). Strikingly, 478 of the 3,000 genes showed a strong UV-induced repositioning of PAF1 to more downstream regions at both 6J/m^2^ and 9J/m^2^ in wild-type cells (Figure 6E; shows pooled data from all replicate experiments of both UV doses), but not in CSB-KO cells (Supplemental Figure 5B). In contrast, 446 genes showed no repositioning of PAF1 upstream of the TSS at both 6J/m^2^ and 9J/m^2^ (Figure 6E). Examples of representative genes are shown in Figure 6F. These findings suggest that the observed repositioning of PAF1 when averaging all 3,000 genes is largely caused by the strong shift in these 478 genes. Notably, the shift of RNAPII into gene bodies after UV irradiation correlated strongly with genes that show PAF1 repositioning (Figure 6H; traveling ratio shown in Supplemental Figure 5D). Indeed, while the 478 genes with strong PAF1 repositioning downstream of the TSS also showed a clear UV-induced release of RNAPII into the gene body, this phenomenon was not observed in the 446 genes that did not show the UV-induced PAF1 repositioning (Fig 6G, H). Importantly, the UV-induced shift in RNAPII binding in these 478 genes was no longer observed at 26 hrs after UV irradiation (Supplemental Figure 5C), suggesting that this reflects increased transcription elongation associated with PAF1 activity that is involved in the recovery of transcription after UV irradiation.

To more directly address if PAF1 is needed for the UV-induced shift of RNAPII, we mapped RNAPII chromatin-binding sites by ChIP-sequencing in OsTIR1 control cells and PAF1-AID knockin cells. To directly compare these conditions, we again calculated the normalized traveling ratio to reflect the extent of RNAPII redistribution in response to UV irradiation in the top 3,000 genes. Importantly, control cells showed a dose-dependent shift of RNAPII after UV irradiation (Figure 6I) similar to what we had observed before (Figure 5J). Conversely, the auxin-induced depletion of PAF1 prevented the UV-induced repositioning of RNAPII (Figure 6I), suggesting that this RNAPII shift is mediated by PAF1. Even when selecting the strongest 902 genes that showed a RNAPII shift after UV in WT cells, we still did not observe this shift in PAF1-depleted cells (Supplemental Figure 5E). The efficient depletion of PAF1 was confirmed by Western blot analysis (Figure 4F), and defective transcription recovery in RRS experiments (Figure 4G, H). Together, these findings show that the repositioning of RNAPII into gene bodies after UV irradiation is dependent on PAF1.

### PAF1 stimulates processive transcription elongation after UV irradiation

During transcription regulation, PAF1C is thought to travel with RNAPII to facilitate efficient elongation through chromatin (Hou et al., 2019; Kim et al., 2010). Indeed, while we could detect PAF1 and RNAPII binding to TTS sites in unirradiated cells, this was substantially reduced at 8 hrs after UV irradiation. Interestingly, PAF1 and RNAPII reads around the TTS site reappeared at 26 hrs after UV (Supplemental Figure S3F, I), suggesting that, upon pause release from promoter-proximal sites, PAF1C may also travel with RNAPII and stimulate processive elongation throughout genes after UV irradiation.

To gain more insight into this possibility, we metabolically pulse-labelled nascent transcripts with bromouridine (BrU) at different time-points after UV irradiation, followed by capture and sequencing of the BrU-labelled nascent RNAs (Figure 7A) (Andrade-Lima et al., 2015). In unirradiated OsTIR1 control cells, transcription reads were evenly distributed between 25 kb and 100 kb of genes. At 3 hrs after UV irradiation, nascent transcription was substantially reduced at TSS sites and progressively decreased further into gene bodies (Figure 7B-D; see replicate in Supplemental Figure 5F) (Andrade-Lima et al., 2015; Williamson et al., 2017). This is consistent with the distribution of DNA lesion in transcribed strands after UV irradiation with 7 J/m^2^ (1 CPD / 16 kb; (Perdiz et al., 2000)), and suggests that these changes reflect the probability of RNAPII molecules encountering a transcription-blocking DNA lesion that limits progression further into the gene. A partial restoration of reads at the TSS and from within gene bodies in OsTIR1 cells could already be detected at 8 hrs after UV, and this was fully restored at 24 hrs after UV (Figure 7B-D). The restoration of nascent transcription near the end of genes was accompanied by the reappearance of both RNAPII and PAF1 at TTS sites detected by ChIP-seq (see the *ZFR* gene in Fig 7D). These data suggest that transcription recovery occurs in a wave starting from the promoter-proximal region and ultimately reaching the end of genes.

**Figure 7.**
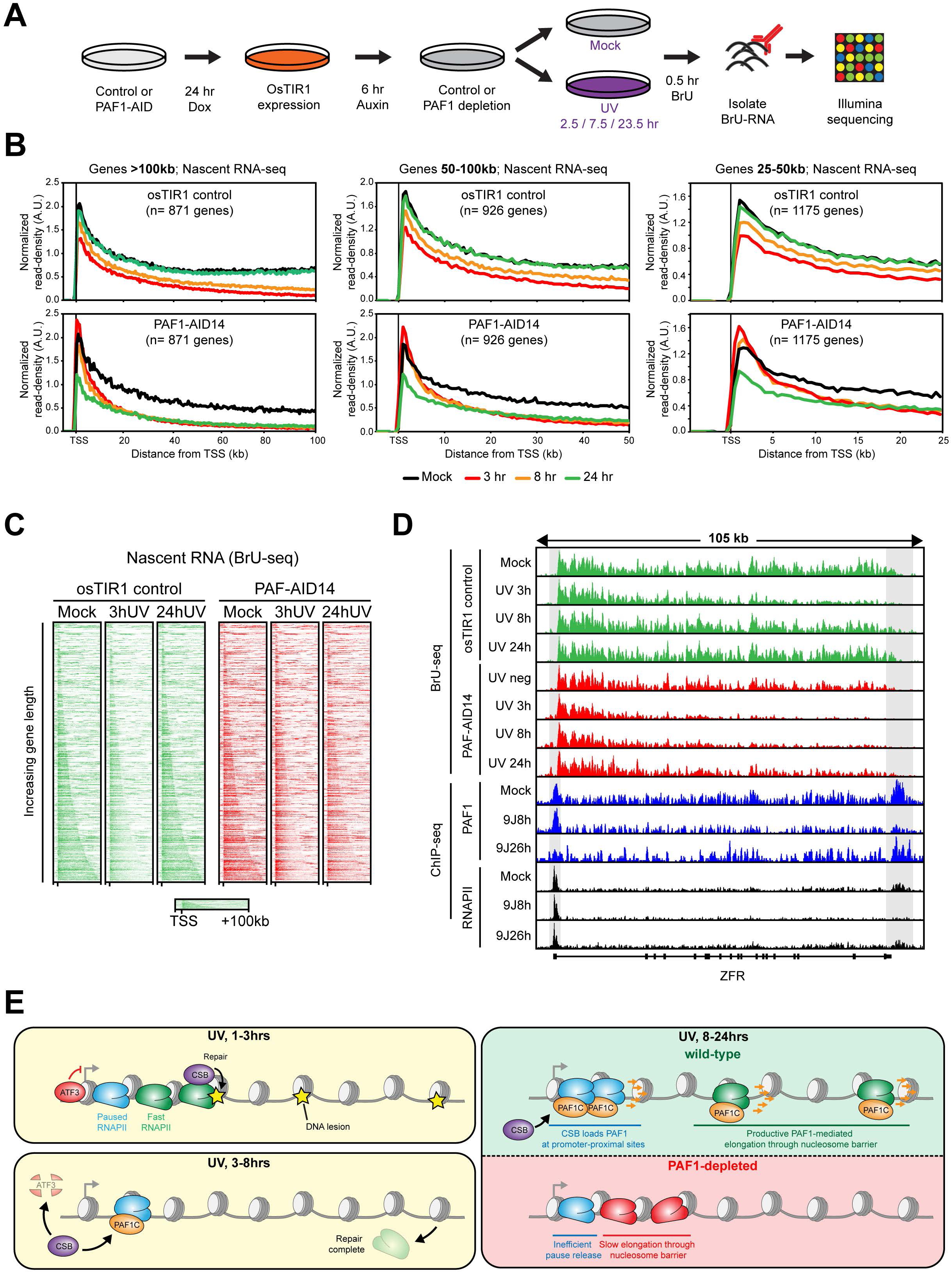
PAF1C activates RNAPII pause release and transcription elongation after UV irradiation. (**A**) Outline of the BrU-seq approach to measure nascent transcription across the genome. (**B**) Metaplots of nascent transcription in genes of >100 kb, between 50 – 100 kb, or between 25 – 50 kb in either osTIR1 cells (upper panels) or PAF1-AID cells (lower panels) that were either mock-treated, or UV-irradiated (7 J/m^2^) and analysed at the indicated time-points (3, 8, or 24 hrs). The relative distribution of nascent transcript read density (in reads per thousand base-pairs per million reads) was normalized to the absolute nascent transcript intensities measured by RRS experiments that were performed in parallel to the BrU-seq experiments using the same cells and time-points (see Figure 4G, H). (**C**) Heatmaps of BrU-seq data from unirradiated (mock) or UV-irradiated (3 or 24 hrs after 7 J/m^2^) osTIR1 control or PAF1-AID cells. Data was mapped and processed as for ChIP-seq and data is presented for the top3000 genes with PAF1 binding at the TSS (as identified in Supplemental Figure 3E) followed by ranking according to gene length. (**D**) UCSC genome browser track showing the nascent transcript read density across the *ZFR* gene in unirradiated and UV-irradiated osTIR1 and PAF1-AID cells. Also shown are the PAF1 and RNAPII read densities for the same gene for comparison. (**E**) Model of the role of PAF1C in restoring transcription across the genome after genotoxic stress.

Unirradiated cells depleted for PAF1 by auxin-mediated degradation showed normal nascent transcription in the first 50 kb and moderately reduced nascent transcription toward the end of long genes (Figure 7B). This is in line with a role of PAF1 as elongation factor (Hou et al., 2019), but also shows that under our experimental conditions there is no dramatic impact of PAF1 depletion on general transcription (Supplemental Figure 5G). Following UV irradiation, PAF1-depleted cells showed a strong and progressive loss of reads into gene bodies at 3 hrs after UV irradiation. However, in contrast to OsTIR1 control cells, PAF1-depleted cells failed to restore RNA synthesis within gene bodies at both 8 and 24 hrs after UV (Figure 7B-D). This impact of PAF1 was most striking for long genes (>100 kb), but also observed in shorter genes (Figure 7B). Nascent transcription in promoter-proximal regions was strongly reduced in OsTIR1 control cells at 3 hrs after irradiation, but fully recovered in a time-dependent fashion. In contrast, transcription in this region in PAF1-depleted cells appeared unaffected at 3 and 8 hrs after UV irradiation, but was strongly reduced at 24 hrs after UV (Figure 7B). Increased nascent transcription around the TSS in PAF1-depleted cells at 8 hrs after UV coincided with increased RNAPII occupancy in this region detected by ChIP-seq (Supplemental Figure 4D) (Hou et al., 2019). Indeed, PAF1-depleted mouse cells were recently shown to accumulate aberrant prematurely terminated transcripts in the TSS region (Hou et al., 2019), which is in line with the increase in nascent transcript species in this region at 3 and 8 hrs after UV (Figure 7B). In contrast, at 24 hrs after UV we find that PAF1-depleted cells show strongly decreased nascent transcription at TSS sites compared to controls (Figure 7B; Supplemental Figure 5G), which is consistent with reduced transcription initiation and / or pause release. Thus, although limited transcription initiation and / or pause release may still be possible without PAF1 at 24 hrs after UV irradiation, we also find that these RNAPII molecules never make it to the end of genes (Figure 7B; Supplemental Figure 5F, G). These results suggest that transcription complexes are not activated for productive and processive elongation throughout genes in the absence of PAF1 after UV irradiation (Figure 7E).

## Discussion

In this study, we define a new CSB-dependent pathway that regulates transcription recovery after TCR by loading the PAF1C elongation complex onto RNAPII. We show that the PAF1C-CSB interaction takes place in promoter-proximal regions where PAF1C triggers RNAPII elongation activation. Our findings support a model in which paused RNAPII molecules at promoter-proximal sites become activated by a CSB-PAF1C axis as a mechanism to restore processive transcription elongation throughout genes following DNA damage repair.

### An alternative CSB pathway links PAF1C to RNAPII for activation

The pTEFb complex containing the CDK9 kinase is involved in the canonical activation of paused RNAPII by triggering the release of NELF and switching DSIF from a negative into a positive elongation factor (Vos et al., 2018b). The activity of CDK9 also triggers the association of the PAF1 complex with RNAPII, resulting in NELF displacement and transcription elongation (Hou et al., 2019; Vos et al., 2018a; Yu et al., 2015). Our interaction data suggests that only a very small RNAPII pool interacts with the PAF1C under steady-state conditions. In fact, we do not detect a robust interaction between these complexes in co-IP or mass spec approaches under our conditions (Figure 1-3). *In vitro* experiments have revealed that the association of PAF1C with RNAPII requires p-TEFb-mediated phosphorylation (Vos et al., 2018a). Thus, the lack of a strong interaction between PAF1C and RNAPII in unirradiated cells could be explained by the presence of only a small pool of p-TEFb-modified RNAPII at any given time.

UV irradiation triggers a strong shift in the pool of RNAPII molecules that engage PAF1C – a process that does not require pTEFb, but is fully dependent on the TCR-specific CSB protein (Figure 3). Since the RNAPII-PAF1C interaction was not affected in other TCR-compromised cells with a knockout for either *CSA*, *UVSSA*, or *XPA* (Figure 3), our findings argue that the loss of the RNAPII-PAF1C interaction in CSB-deficient cells is not a general result of TCR deficiency or associated with the decrease in the hypo-phosphorylated form of RNAPII after UV (Rockx et al., 2000). It is possible that CSB mediates the association of PAF1C to RNAPII through protein-protein interactions. Although speculative, the cryo-EM structures of the yeast CSB orthologue, RAD26 (Xu et al., 2017), and the human PAFC1 complex bound to RNAPII (Vos et al., 2018a) are, in principle, compatible with direct contacts between these complexes (Supplemental Figure 6).

During the canonical transition of paused RNAPII into productive elongation, the association of PAF1C displaces the NELF complex from RNAPII. Interestingly, a p38 MAP kinase pathway releases NELF from chromatin after UV irradiation, which is partially dependent on CSB (Borisova et al., 2018). Importantly, the UV-induced association of PAF1C with RNAPII still occurred following inhibition of p38 signalling (Figure 5A), suggesting that CSB-dependent PAF1 recruitment does not occur downstream from p38-mediated NELF release. In light of these findings, we favour a scenario in which CSB could promote efficient NELF displacement by both regulating PAF1C recruitment and activating the p38 pathway in parallel.

In addition to NELF release, p38 also activates a p-TEFb pathway that regulates a transcriptional response immediately after DNA damage induction by ensuring the expression of short coding and non-coding RNAs involved in the DNA damage response, including *FOS* and *CDKN1A* (Bugai et al., 2019). Importantly, many of these short DDR genes are regulated by p53 and do not require CSB for their expression after UV irradiation (Proietti-De-Santis et al., 2006). Thus, it appears that cells mount an immediate transcriptional response through p-TEFb to ensure expression of short DDR genes and non-coding RNAs while most other genes undergo a transcriptional arrest, which is particularly striking for longer genes (Andrade-Lima et al., 2015; Bugai et al., 2019; Williamson et al., 2017). We here show that the transcription recovery of those arrested genes following repair by TCR requires the CSB-PAF1C axis for efficient recovery of productive elongation.

### A dual role for CSB in DNA repair and transcription recovery

Our findings suggest a dual role for CSB in regulating transcription recovery after UV irradiation. Firstly, CSB is an essential DNA repair factor in TCR that associates with DNA damage-stalled RNAPII and subsequently facilitates the recruitment of downstream TCR factors to initiate repair (unpublished data). Secondly, CSB has another role that is independent of repair in regulating transcription recovery from promoter-proximal sites, which involves the CSB-mediated association of PAF1C with RNAPII (Figure 1-3). We show that PAF1C is dispensable for the repair of transcription-blocking DNA lesions, suggesting that the PAF1C-CSB interaction plays a unique role in transcription recovery. Moreover, several lines of evidence suggest that the CSB–PAF1C interaction may take place at promoter-proximal regions. Firstly, we find that transcription initiation inhibitors indeed strongly affect the PAF1C-RNAPII interaction, while specific transcription elongation inhibitors do not (Figure 5A). Secondly, PAF1 binding and nascent transcription (Supplemental Figure 3G; Figure 7) became restricted to promoter-proximal regions within the first hours after UV, suggesting that this is the region where the UV-induced PAF1-RNAPII interaction takes place. Strikingly, CSB is already bound near TSS sites of genes that also bind PAF1 and RNAPII (Figure 6) (Epanchintsev et al., 2017), which could reflect its role in regulating transcription elongation (Selby and Sancar, 1997).

Our interaction studies reveal that PAF1C in unirradiated cells does not associate strongly with either CSB or RNAPII (Figure 2, Supplemental Figure 1), so why do these proteins seem to co-localize in the genome? There are several ways to explain these data. Firstly, we use protein-protein and DNA-protein crosslinking in the ChIP-seq experiments, which could stabilize even transient interactions between proteins, while our mass spec interaction studies are performed under stringent conditions without crosslinking. It is possible that CSB and PAF1C transiently interact in unirradiated cells, but that this interaction is strongly stabilized after UV, explaining why we detect such as robust UV-induced association between CSB-PAF1C-RNAPII after UV. Secondly, it is possible that these proteins are independently bound to the same genomic regions without interacting at the protein level. In line with this interpretation, we find that RNAPII chromatin-bound levels at TSS sites drastically drop in CSB-deficient cells after UV irradiation, while the PAF1 bound levels at TSS sites remain unchanged (Figure 6). These findings argue that PAF1 is bound to these sites independently of RNAPII and CSB. Indeed, recent studies in yeast have shown that PAF1C binds directly to nucleosomes through the acidic patch (Cucinotta et al., 2019), suggesting that this is certainly possible.

### PAF1-mediated activation of transcription elongation from promoter-proximal sites

At 8 hours after UV irradiation, we detect a marked shift in the read density of both PAF1 and RNAPII promoter-proximal peaks into the first ∼2 kb of gene bodies (Figure 5-6). These changes could be triggered by the strong UV-induced interaction between PAF1C and RNAPII, which is facilitated by CSB. One interpretation is that these changes reflect increased elongation from promoter-proximal sites, resulting in an increased number of RNAPII molecules within this region (Figure 5-6). Alternatively, this increase in read density could also be due to decreased processivity of RNAPII and bound PAF1 causing their accumulation in promoter-proximal regions (Ehrensberger et al., 2013; Hou et al., 2019). Although both options are possible and fully compatible with our model, we favour the interpretation that PAF1 and RNAPII repositioning at the 5’-end of genes reflect increased elongation. In line with this, we find that genes that do not show a shift in PAF1 binding also do not release RNAPII, and depletion of PAF1 prevents the UV-induced release of RNAPII from TSS sites (Figure 6).

While the binding profile of CSB around TSS sites is unaltered by UV irradiation, CSA becomes strongly bound in these regions after UV (Figure 6) (Epanchintsev et al., 2017). These findings suggest the following scenario (Figure 7E): i) CSB is already bound around TSS sites in unirradiated cells, ii) these CSB molecules recruit PAF1C after UV and load this complex onto RNAPII paused at these sites, iii) CSB recruits CSA to TSS sites after UV irradiation resulting in the ubiquitylation of ATF3, which is also bound near the TSS sites to repress transcription (Epanchintsev et al., 2017), iv) the release of the repressive impact of ATF3 combined with the activation of RNAPII by PAF1C promotes pause release and elongation activation that drives transcription recovery. Our findings also suggest that PAF1 that is loaded onto RNAPII by CSB at promoter-proximal sites travels with RNAPII and facilitates efficient and productive elongation throughout the gene. Supporting this notion, we find that PAF1 at TTS sites is strongly reduced shortly after UV, but reappears at 26 hrs after UV irradiation (Figure 7D; Supplemental Figure 3F, I). At this time, we also detect that RNA synthesis is restored near the end of genes in wild-type cells, but not in PAF1-depleted cells (Figure 7). In this model, PAF1C stimulates productive transcription elongation throughout gene bodies (Figure 7E), which is consistent with recently published data (Hou et al., 2019). An intriguing possibility is that PAF1C is needed to stimulate transcription through former repair sites, which may have a chromatin signature that is suboptimal for transcription, such as H2A ubiquitylation (Bergink et al., 2006).

## Experimental Procedures

### Cell lines

All cell lines are listed in Table 1. All human RPE-1-hTERT-Flp-In/T-Rex, U2OS, U2OS-Flp-In/T-REx and CSB-deficient CS1AN-SV cells were cultured at 37°C in an atmosphere of 5% CO_2_ in DMEM, supplemented with antibiotics, 10% fetal calf serum and glutaMAX (Gibco). Primary XP-C patient fibroblasts XP168LV were cultured at 37°C in an atmosphere of 5% CO_2_ in Ham’s F10 medium without thymidine (Lonza) supplemented with 20% fetal calf serum and antibiotics.

**Table 1.** Cell lines used in this study.

| Target | Source |
| --- | --- |
| CS1AN-SV40 + GFP | LUMC, dr. L. Mullenders |
| CS1AN-SV40 + GFP-CSB | LUMC, dr. L. Mullenders |
| RPE-hTERT(FRT) | Ximbio, London, UK |
| RPE-hTERT(FRT) + GFP-LEO1 clone 6 | This study |
| RPE-hTERT(FRT) + GFP-NLS | This study |
| U2OS GFP-RPB1 clone I | LUMC, dr. H. van Attikum (Caron et al., 2019) |
| U2OS GFP-RPB1 CSA-KO clone 2-4 | This study |
| U2OS GFP-RPB1 CSB-KO clone 1-40 | This study |
| U2OS GFP-RPB1 UVSSA-KO clone 1-4 | This study |
| U2OS TetOn-OsTIR1 | LUMC, dr. H. van Attikum |
| U2OS TetOn-OsTIR1 PAF-AID clone 14 | This study |
| U2OS TetOn-OsTIR1 PAF-AID clone 15 | This study |
| U2OS(FRT) | This study |
| U2OS(FRT) + siRNA-resistant GFP-PAF1 | This study |
| U2OS(FRT) CSA-KO + CSA-GFP (dox) clone X | This study |
| U2OS(FRT) CSA-KO clone 2-16 | This study |
| U2OS(FRT) CSB-KO + GFP-CSB (dox) clone 3 | This study |
| U2OS(FRT) CSB-KO clone 1-12 | This study |
| U2OS(FRT) UVSSA-KO + UVSSA-GFP (dox) clone 1-3 | This study |
| U2OS(FRT) UVSSA-KO clone 1-8 | This study |
| U2OS(FRT) XPA-KO + GFP-XPA (dox) clone 8 | This study |
| U2OS(FRT) XPA-KO clone 2-8 | This study |
| U2OS(FRT) XPC-KO clone 2-7 | This study |
| XP168LV primary fibroblasts | Erasmus MC; dr. J. Marteijn (Wienholz et al., 2017) |

Flp-In/T-REx cells (either RPE1-hTERT or U2OS) were used to stably express inducible version of GFP-tagged proteins by co-transfecting pCDNA5/FRT/TO-Puro plasmid encoding GFP-tagged fusion proteins (5 µg), together with pOG44 plasmid encoding the Flp recombinase (0.5 µg). After selection on 1 µg/mL puromycin, single clones were isolated and expanded. RPE-1-hTERT-Flp-In/T-Rex were generated expressing either GFP-NLS or GFP-LEO1. U2OS-Flp-In/T-Rex (knock-out for specific TCR genes; see below) were generated stably expressing CSA-GFP, GFP-CSB, UVSSA-GFP, or GFP-XPA in the corresponding KO line. In addition, U2OS-Flp-In/T-Rex were generated expressing siRNA-resistant PAF1. Stable U2OS-Flp-In/T-REx or RPE-hTERT-U2OS-Flp-In/T-REx clones were incubated with 2 µg/mL doxycycline to induce expression of GFP-tagged proteins.

To generate cells sensitive to auxin-inducible degradation of PAF1, U2OS cells expressing OsTIR1 under the control of doxycycline (U2OS-TetOn-OsTIR1) were transfected with plasmids encoding Cas9 and an sgRNA close to the stop codon of the PAF1 gene, together with a donor plasmid containing an auxin-inducible degron (AID) and G418 cassette (AID-P2A-G418) flanked by ∼1 kb arms homologous to the PAF1 locus (Supplemental Figure 3A). This generated endogenously-tagged U2OS-TetOn-OsTIR1-PAF-AID cells. Cells were selected with 200 µg/mL G418 for ∼14 days and individual clones were selected and tested for auxin-inducible PAF1 degradation using western blot analysis. To induce depletion of PAF1, cells were induced to express OsTIR1 by ∼24 hrs treatment with 2 µg/mL Doxycycline, followed by treatment with 500 µM auxin (3-Indoleacetic acid; Sigma) for 5-6 hrs.

### Generation of TCR knock-out cells

U2OS-Flp-In/T-Rex were co-transfected with pU6-gRNA:PGK-puro-2A-tagBFP (Sigma-Aldrich library from the LUMC) containing specific sgRNA targeting *CSB*, *CSA*, *UVSSA*, or *XPA* (listed in Table 2), together with pX458 (addgene) encoding Cas9. Cells were selected with puromycin (1 µg/ml) for 3 days and seeded at low density without puromycin. Individual clones were isolated and screened for loss of protein-of-interest expression and absence of stable Cas9 expression by western blot analysis and/ or sanger sequencing.

**Table 2.**
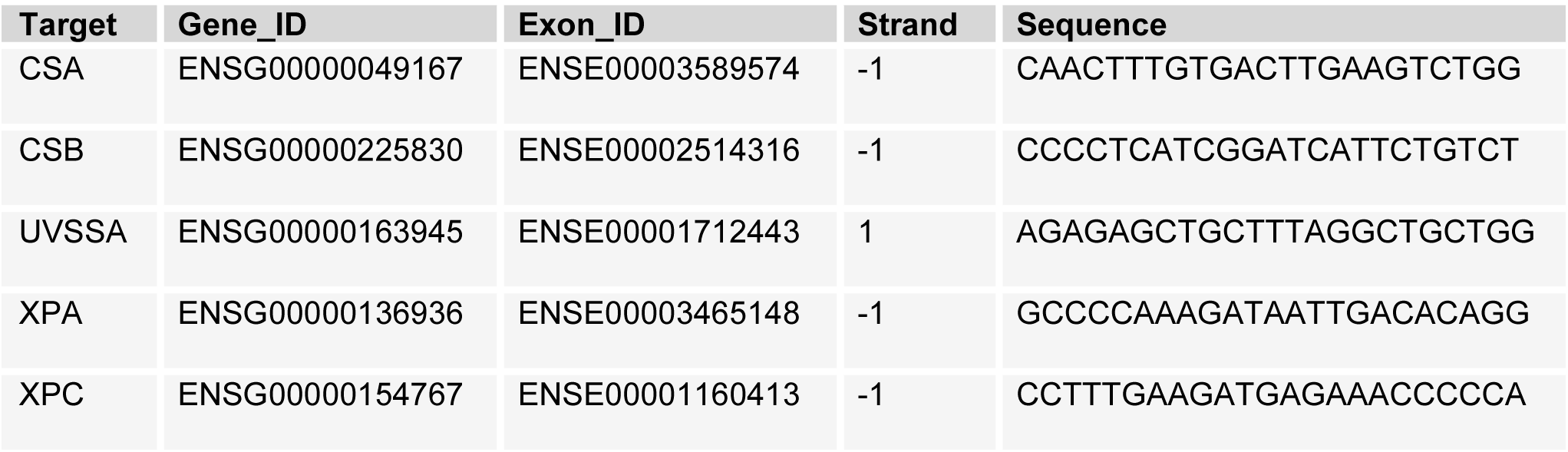
sgRNAs used in this study.

### Plasmids

pcDNA5/FRT/TO-Puro was purchased from Addgene. PCR was used to generate the following GFP fusion proteins, which were inserted into pcDNA5/FRT/TO-Puro: GFP-NLS, GFP-CSB, CSA-GFP, UVSSA-GFP, GFP-XPA, GFP-LEO1, GFP-CTR9, GFP-PAF1. Overlap PCR was used to generate GFP-PAF1 that was resistant to siPAF1-2 and siPAF1-3 by introducing the following silent mutations: 5-AA<u>A</u> CA<u>A</u> CA<u>A</u> TT<u>C</u> AC<u>A</u> GA<u>A</u> GA<u>G</u>-3 and 5-GA<u>C</u> GA<u>C</u> GT<u>C</u> TA<u>C</u> GA<u>T</u> TA<u>T</u>-3.

### Transfections

Cells were transfected with plasmid DNA using Lipofectamine 2000 according to the manufacturer’s instructions. Cells were typically imaged 24 hrs after transfection. All siRNA transfections (see list of siRNA sequences in Table 3) were performed with 40 nM siRNA duplexes using Lipofectamine RNAiMAX (Invitrogen). Cells were transfected twice with siRNAs at 0 and 36 hrs and were typically analyzed 60 hrs after the first transfection.

**Table 3.**
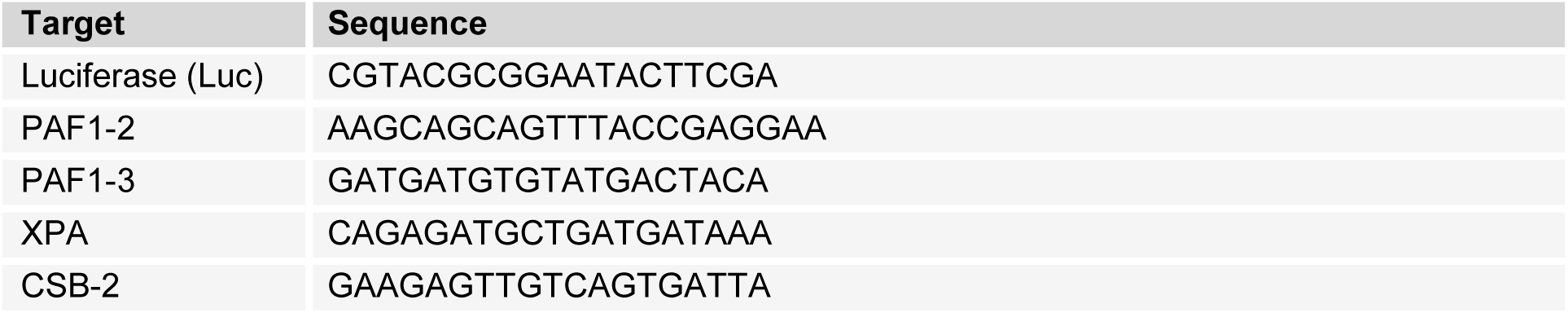
siRNAs used in this study.

### Western blotting

Cell extracts were generated by cell lysis and boiled in sample buffer. Proteins were separated by sodium dodecyl sulfate polyacrylamide gel electrophoresis (SDS-PAGE) and transferred to nitrocellulose membranes. Protein expression was analyzed by immunoblotting with the indicated primary antibodies (listed in Table 4) and secondary CF680 Goat Anti-Rabbit IgG antibody at 1:10,000, CF770 Goat Anti-Mouse IgG antibody at 1:10,000, followed by detection using the Odyssey infrared imaging scanning system (LI-COR biosciences, Lincoln, Nebraska USA).

**Table 4.** Antibodies used in this study.

| Antibody | Host | Company (reference) | Use | Antibody identifier |
| --- | --- | --- | --- | --- |
| Cas9 | Mouse | Cell signalling (7A9-3A3 #14697) | WB: 1:2,000 | aML#031 |
| CHD4 | Rabbit | Active Motif (39289) | WB: 1:1,000 | aML#019 |
| CSA | Rabbit | Abcam (ab137033) | WB: 1:750 | aML#028 |
| CSA | Mouse | Santa Cruz (sc376981) | WB: 1:500 | aML#025 |
| CSB | Rabbit | Santa Cruz (sc25370) | WB; 1:200 | aML#003 |
| CSB | Goat | Santa Cruz (SC-10459, E-18) | WB; 1:1,000 | aML#039 |
| CTR9 | Rabbit | Bethyl (A301-395A) | WB; 1:5,000 | aML#049 |
| GFP | Mouse | Roche (11814460001) | WB; 1:1,000 | aML#011 |
| GFP | Rabbit | Abcam (ab290) | WB; 1:1,000 | aML#044 |
| LEO1 | Rabbit | Bethyl, (A300-175A) | WB; 1:3,000 | aML#023 |
| PAF1 | Rabbit | Bethyl (A300-172A) | WB; 1:3,000<br>ChIP; 3µg | aML#022 |
| RPB1 (total) | Rabbit | Bethyl (A304-405A) | ChIP: 3µg | aML#088 |
| RPB1-S2 | Rabbit | Abcam (ab5095) | WB; 1:1,000 | aML#024 |
| RPB1-S5 | Mouse | Abcam (ab5408) | WB: 1:1,000 | - |
| Tubulin | Mouse | Sigma (T6199) | WB; 1:1,000 | aML#008 |
| XPA | Mouse | Invitrogen (MA5-13835) | WB; 1:500 | aML#002 |
| XPA | Rabbit | gift of Rick Wood (CJ1) | WB; 1:10,000 | aML#079 |
| XPC | Mouse | Abcam (ab6264) | WB: 1:500 | aML#001 |

### Clonogenic Survival assays

Cells were plated in low density in culture dishes, allowed to attach and treated with Illudin S in different concentrations for 72 hrs. Illudin S was removed and cells were allowed to form clones for 7-10 days. To visualize clones, cells were subjected to NaCl fixation and methylene blue staining. Cell survival after Illudin S treatment was defined as the percentage of cells able to form clones, relative to the untreated condition.

### Immunoprecipitation for Co-IP

Except where indicated otherwise, all co-IP experiments were performed 1h after UV-irradiation. For endogenous RNAPII immunoprecipitation, cells were subjected to chromatin fractionation prior to immunoprecipitation. Cells were lysed in EBC-150 buffer (50 mM Tris, pH 7.5, 150 mM NaCl, 0.5% NP-40, 2 mM MgCl_2_ supplemented with protease and phosphatase inhibitor cocktails (Roche)) for 20 minutes at 4°C, followed by centrifugation to remove cytoplasmic proteins. Subsequently, the chromatin fraction was solubilized in EBC-150 buffer with 500 U/mL Benzonase and 2 ug of antibody directed to endogenous RNAPII-S2 (Abcam; ab5095) or RNAPII-S5 (Abcam; ab5408) for 1 hr at 4°C under rotation. Next, the NaCl concentration of the lysis buffer was increased to 300 mM by adding concentrated (5M) NaCl, and lysates were incubated for another 30 min at 4°C. The lysates were cleared from insoluble chromatin and were subjected to immunoprecipitation with protein A agarose beads (Millipore) for 1.5 hrs at 4°C. The beads were then washed 4-6 times with EBC-300 buffer (50 mM Tris, pH 7.5, 300 mM NaCl, 0.5% NP-40, 1 mM EDTA) and boiled in sample buffer. Bound proteins were resolved by SDS-PAGE and immunoblotted with the indicated antibodies.

Immunoprecipitation of GFP-tagged proteins was performed using a similar protocol with the following exceptions: Cell pellets were directly solubilized in EBC-150 supplemented with 500U Benzonase, without chromatin fractionation, and pulldown was not dependent on antibody-mediated pulldown, but was performed using GFP Trap beads (Chromotek).

### Generation of mass spectrometry samples

For stable isotope labeling by amino acids in cell culture (SILAC) labeling, CS1AN-SV expressing GFP-CSB were cultured for 14 days in media containing ‘heavy’ (H) and ‘light’ (L) labeled forms of the amino acids arginine and lysine respectively. SILAC-labeled cells were mock-treated (L) or exposed to 20 J/m^2^ UV-C light and allowed to recover for 1 hr (H). Label-free mass spectrometry samples were also either kept untreated or exposed to 20 J/m^2^ UV-C light and allowed to recover for 1 hr. A pool of equal amounts of H-and L-labelled cells (SILAC) or individual label-free mass spectrometry samples, were subsequently subjected to immunoprecipitation using GFP Trap beads as described above. After pulldown, the beads were washed 2 times with EBC-300 buffer and 2 times with 50 mM (NH_4_)_2_CO_3_ followed by overnight digestion using 2.5 µg trypsin at 37°C under constant shaking. Peptides of the H and L precipitates were mixed in a 1:1 ratio and all samples were desalted using a Sep-Pak tC18 cartridge by washing with 0.1 % acetic acid. Finally, peptides were eluted with 0.1 % formic acid / 60 % acetonitrile and lyophilized according to (Rappsilber et al., 2007).

### Mass spectrometry data acquisition

Mass spectrometry was performed essentially as previously described (Kumar et al., 2017). Samples were analyzed on a Q-Exactive Orbitrap mass spectrometer (Thermo Scientific, Germany) coupled to an EASY-nanoLC 1000 system (Proxeon, Odense, Denmark). For the SILAC samples, digested peptides were separated using a 13 cm fused silica capillary (ID: 75 µm, OD: 375 µm, Polymicro Technologies, California, US) in-house packed with 1.8 µm C18 beads (Reprospher-DE, Pur, Dr. Maisch, Ammerburch-Entringen, Germany). Peptides were separated by liquid chromatography using a gradient from 2% to 95% acetonitrile with 0.1% formic acid at a flow rate of 200 nl/min for 2 hrs. The mass spectrometer was operated in positive-ion mode at 1.8 kV with the capillary heated to 250°C. Data-dependent acquisition mode was used to automatically switch between full scan MS and MS/MS scans, employing a top 10 method. Full scan MS spectra were obtained with a resolution of 70,000, a target value of 3×10^6^ and a scan range from 400 to 2,000 m/z. Higher-Collisional Dissociation (HCD) tandem mass spectra (MS/MS) were recorded with a resolution of 17,500, a target value of 1×10^5^ and a normalized collision energy of 25%. Maximum injection times for MS and MS/MS were 20ms and 60ms, respectively. For Label free samples, digested peptides were separated using a 15 cm fused silica capillary (ID: 75 µm, OD: 375 µm, Polymicro Technologies, California, US) in-house packed with 1.9 µm C18-AQ beads (Reprospher-DE, Pur, Dr. Maisch, Ammerburch-Entringen, Germany). Peptides were separated by liquid chromatography using a gradient from 2% to 95% acetonitrile with 0.1% formic acid at a flow rate of 200 nl/min for 90 min. The mass spectrometer was operated in positive-ion mode at 2.8 kV with the capillary heated to 250°C. Data-dependent acquisition mode was used to automatically switch between full scan MS and MS/MS scans, employing a top 7 method. Full scan MS spectra were obtained with a resolution of 70,000, a target value of 3×10^6^ and a scan range from 400 to 2,000 m/z. Higher-Collisional Dissociation (HCD) tandem mass spectra (MS/MS) were recorded with a resolution of 35,000, a target value of 1×10^5^ and a normalized collision energy of 25%. Maximum injection times for MS and MS/MS were 50ms and 120ms, respectively. For all samples, the precursor ion masses selected for MS/MS analysis were subsequently dynamically excluded from MS/MS analysis for 60 sec. Precursor ions with a charge state of 1 or greater than 6 were excluded from triggering MS/MS events.

### Mass spectrometry data analysis

Raw mass spectrometry data were further analysed in MaxQuant v 1.5.3.30 according to (Tyanova et al., 2016a) using standard settings with the following modifications, For the SILAC-labelled GFP-CSB samples, multiplicity was set to 2, marking Arg10 and Lys8 as heavy labels. Maximum missed cleavages by trypsin was set to 4. Searches were performed against an *in silico* digested database from the human proteome including isoforms and canonical proteins (Uniprot, 18^th^ June 2018). Minimum peptide length was set to 6 aa and maximum peptide mass was set to 5 kDa. Carbamidomethyl (C) was disabled as fixed modification. The match between runs feature was activated. Minimum ratio count for quantification was set to 1. For the label-free GFP-LEO1 and GFP-RBP1 samples, maximum missed cleavages by trypsin was set to 4. Label-free quantification was activated, not enabling Fast LFQ. Searches were performed against an *in silico* digested database from the human proteome including isoforms and canonical proteins (Uniprot, 18^th^ June 2018). Carbamidomethyl (C) was disabled as fixed modification. The match between runs feature was activated and iBAQ quantification was also enabled. MaxQuant output data from the SILAC samples analysis were further processed in Microsoft Excel 2016 for comprehensive visualization. Label-free analysis was further carried out in the Perseus Computational Platform v1.5.5.3 according to (Tyanova et al., 2016b). LFQ intensity values were log2 transformed and potential contaminants and proteins identified by site only or reverse peptide were removed. Samples were grouped in experimental categories and proteins not identified in 4 out of 4 replicates in at least one group were also removed. Missing values were imputed using normally distributed values with a 1.8 downshift (log2) and a randomized 0.3 width (log2) considering whole matrix values. Two sample t-tests were performed to compare groups. Analyzed data were exported from Perseus and further processed in Microsoft Excel 2016 for comprehensive visualization. The mass spectrometry proteomics data have been deposited to the ProteomeXchange Consortium via the PRIDE partner repository with the dataset identifier PXD016198 (Perez-Riverol et al., 2019).

### RNA recovery synthesis (RRS) assay

Cells were plated in DMEM supplemented with 10% Fetal Calf Serum (FCS) and, if needed, transfected with siRNAs as described above. Subsequently, cells were placed in DMEM supplemented with 1% FCS for at least 24 hrs prior to the RRS experiment to reduce the excess of available uridine in the culture medium. Cells were UV irradiated, allowed to recover for the indicated time periods, and pulse-labelled with 400 µM 5-ethynyl-uridine (EU; Axxora) for 1 hr. After medium-chase with DMEM without supplements for 15 min, cells were fixed with 3.7% formaldehyde in phosphate-buffered saline (PBS) for 15 min and stored in PBS. Nascent RNA was visualized by click-it chemistry, labeling the cells for 1 hr with a mix of 60 µM atto azide-Alexa594 (Atto Tec), 4 mM copper sulfate (Sigma), 10 mM ascorbic acid (Sigma) and 0.1 μg/mL DAPI in a 50 mM Tris-buffer. Cells were washed extensively with PBS and mounted in Polymount (Brunschwig).

### TCR-specific unscheduled DNA synthesis

Detection of TCR-specific unscheduled DNA synthesis was performed essentially as previously described (Wienholz et al., 2017). Primary XP168LV (XP-C patient cells) were transfected with siRNAs and subsequently serum starved for at least 24 hrs in F10 medium (Lonza) supplemented with 0.5% FCS and antibiotics. Cells with subsequently irradiated with 8 J/m^2^ UV-C, and pulse-labelled with 20 µM 5-ethynyl deoxy-uridine (EdU; Invitrogen) and 1 µM FuDR (Sigma Aldrich) for 8 hrs. After labelling, cells were chased with F10 medium supplemented with 0.5% FCS and 10 µM thymidine for 15 minutes, and fixed for 15 min with 3.6% formaldehyde and 0.5% Triton-X100 in PBS. Next, cells were permeabilized for 20 min in PBS with 0.5% Triton-X100 and washed and stored in 3% bovine serum albumin (BSA, Thermo Sisher) in PBS. The incorporated EdU was visualized by click-it chemistry-mediated binding of Biotin (Azide-PEG3-Biotin Conjugate; Jena Biosciences) using the protocol and reagents from the Invitrogen Click-iT EdU Cell Proliferation Kit for Imaging (Invitrogen), and signals were amplified using protocol and reagent of the Alexa Fluor-488 Tyramide streptavidin SuperBoos Kit (Thermo Scientific). After click-it and amplification, cells were counterstained with 0.1 μg/mL DAPI, washed extensively with 0.1% Triton-X100 in PBS and mounted in Polymount (Brunschwig).

### Microscopic analysis of fixed cells

Images of fixed samples were acquired on a Zeiss AxioImager M2 or D2 widefield fluorescence microscope equipped with 63x PLAN APO (1.4 NA) oil-immersion objectives (Zeiss) and an HXP 120 metal-halide lamp used for excitation. Fluorescent probes were detected using the following filters: DAPI (excitation filter: 350/50 nm, dichroic mirror: 400 nm, emission filter: 460/50 nm), Alexa 555 (excitation filter: 545/25 nm, dichroic mirror: 565 nm, emission filter: 605/70 nm), Alexa 647 (excitation filter: 640/30 nm, dichroic mirror: 660 nm, emission filter: 690/50 nm). Images were recorded using ZEN 2012 software and analyzed in Image J.

### ChIP-sequencing

Cells were plated and grown to ∼90% confluency and crosslinked with 0.5 mg/mL disuccinimidyl glutarate (DSG; Thermo Fisher) in PBS for 45 min at room temperature. Cells were washed with PBS and crosslinked with 1 % PFA for 20 min at room temperature. Fixation was stopped by adding 1.25 M Glycin in PBS to a final concentration of 0.1 M for 3 minutes at room temperature. Cells were washed with cold PBS and lysed and collected in a buffer containing 0.25% Triton X-100, 10 mM EDTA (pH 8.0), 0.5 mM EGTA (pH 8.0) and 20 mM Hepes (pH 7.6). Chromatin was pelleted in 5 min at 400 g and incubated in a buffer containing 150 mM NaCl, 1 mM EDTA (pH 8.0), 0.5 mM EGTA (pH 8.0) and 50 mM Hepes (pH 7.6) for 10 minutes at 4°C. Chromatin was again pelleted for 5 min at 400 g and resuspended in ChIP-buffer (0.15 % SDS, 1 % Triton X-100, 150 mM NaCl, 1 mM EDTA (pH 8.0), 0.5 mM EGTA (pH 8.0) and 20 mM Hepes (pH 7.6)) to a final concentration of 15×10^6^ cells/ml. Chromatin was sonicated to approximately 1 nucleosome using a Bioruptor waterbath sonicator (Diagenode). Chromatin of ∼5×10^6^ cells was incubated with 3ug antibody (RNAPII, rabbit polyclonal, Bethyl laboratories, A304-405A; PAF1, rabbit polyclonal, Bethyl laboratories, A300-172A) over night at 4°C, followed by a 1.5 hrs protein-chromatin pull-down with a 1:1 mix of protein A and protein G Dynabeads (Thermo Fisher; 10001D and 10003D). ChIP samples were washed extensively, followed by decrosslinking for 4 hrs at 65°C in the presence of proteinase K. DNA was purified using a Qiagen MinElute kit. Sample libraries were prepared using Hifi Kapa sample prep kit and A-T mediated ligation of Nextflex adapters. Samples were sequenced using an Illumina NextSeq500, using paired-end sequencing with 42 bp from each end. Raw data and processed files are deposited in the Gene Expression Omnibus (GEO).

### ChIP-seq analyses

Reads were aligned to the Human Genome 38 (Hg38) using bwa-mem tools (Li, 2013). Only high-quality reads (> q30) were included in the analyses and duplicates were removed. Bedgraph UCSC genome tracks were generated and PAF1 binding peaks were identified using the callpeaks tool of MACS2 (Zhang et al., 2008), loading sample versus UV-dose-associated input with standard tool-settings. Bam files were converted into TagDirectories using HOMER tools (Heinz et al., 2010). Binding profiles within selected areas of individual genes (e.g. around TSS or TTS), were defined using the AnnotatePeaks.pl tool of HOMER using the default normalization to 10mln reads. Metagene profiles were defined using the makeMetaGeneProfile.pl tool of HOMER, using default settings. Individual datasets were subsequently processed in R (Team, 2019). First, read densities in input samples were subtracted from individual ChIP-seq datasets to background-correct our data in which negative values were converted to 0, to prevent the use of impossible negative read densities in further calculations. ChIP profiles were averaged per sets of genes and profiles were normalized to area under the curve to allow proper comparison of the profiles without effects of overall differences in read density. Traveling ratios were defined per gene over ranges indicated in individual analyses, with infinite ratios removed. A list of 49,948 transcription start sites was obtained from the UCSC genome database (https://genome.ucsc.edu/cgi-bin/hgTables) selecting the knownCanonical table containing the canonical transcription start sites per gene. Only genes of 3-100kb were included in the analyses to prevent the inclusion of extremely small genes that might not be damaged under our experimental conditions, or genes that likely will acquire multiple DNA damages and might therefore not represent repaired genes in the timeframe of our experiments. To prevent contamination of binding profiles, genes should be non-overlapping with at least 2 kb between genes. A total of 8,811 genes were selected. Original ChIP-seq datafiles for RNAPII, ATF3, CSA and CSB, were also obtained from Epanchintsev and colleagues (Epanchintsev et al., 2017) (GSE87562), and data for PAF1 was obtained from Chen and colleagues (Chen et al., 2017) (GSE97527). These files were subsequently converted into FASTQ files using NCBI sratoolkit.2.9.6-1-win64 and processed as described above, but without subtraction of Input reads.

### BrU-sequencing and data analysis

U2OS osTIR1 or PAF1-AID cells were induced with doxycycline for 24 hrs, and subsequently with auxin for 5 hrs. After this treatment, cells were either mock-treated or irradiated with UV-C light (7 J/m^2^). Cells were then incubated in conditioned media for different periods of time (0, 3, 8, 24h) before being incubated with 2 mM bromouridine (Bru) at 37°C for a 30 min. The cells were then lysed in TRIzol reagent (Invitrogen) and Bru-containing RNA isolated as previously described (Andrade-Lima et al., 2015). cDNA libraries were made from the Bru-labeled RNA using the Illumina TruSeq library kit and paired-end 151 bp sequenced using the Illumina NovaSeq platform at the University of Michigan DNA Sequencing Core. Data was processed as previously described (Paulsen et al., 2014). Briefly, reads were pre-filtered by alignment to the human ribosomal repeating subunit (GenBank U13369.1) and human mitochondrial genome (chrM) from the hg38 reference genome using bowtie2 v.2.3.3.1. The remaining reads were mapped to the hg38 reference genome using STAR v2.7.0f. Base coverages were used to compute read counts for features, such as genes and bins, which were then normalized to feature length and number of uniquely-mapped reads (RPKM method). Gene selection (n = 871) was based on the following criteria: length of at least 100kb, TSS is at least 10kb apart, expression is at least 0.05 RPKM. The median expression was calculated for each 500 bp bins from 5kb upstream until 100kb downstream of the selected genes for each time-point. Aggregate plots were normalized to total RNA levels, as quantified by 5-EU labelling (RRS), relative to the control in their specific cell type. For example, PAF-AID cells 3 hours after UV irradiation showed 62% RNA relative to PAF-AID control cells, so we multiplied the expression of the bins by 0.62. Heatmaps and UCSC tracks were generated by mapping and processing data as described for ChIP-seq analyses. Raw data and processed files are deposited in the Gene Expression Omnibus (GEO).

## Author Contributions

DvdH generated knockout cells, constructs and stable cell-lines, PAF1-AID knockin cells, performed clonogenic survivals, PCR and Western blot analysis to validate knockouts, RRS experiments, Co-IP experiments for Western blot analysis, Co-IP experiments for mass spectrometry, ChIP experiments and analysis, BrU pulse-labelling experiments and analysis, and wrote the paper. CGS performed ChIP-seq experiments, and measured and analyzed MS experiments in Figure 1B. MV supervised CGS and provided the infrastructure for ChIP-seq experiments. RG-P and ACOV analyzed all MS experiments except those in Figure 1B. AK and YvdW generated knockout cells, stable cell-lines, and performed RRS experiments. KA generated knockout cells and stable cell-lines. MD performed SILAC-based MS. MTP captured nascent RNA and performed library preparations for the Bru-seq experiments. KY and ML analyzed the Bru-seq data. DZ and JAFM performed and analyzed TCR-UDS experiments. HW and LD provided tools for ChIP-seq analyses. MSL conceived and supervised the project and wrote the paper.

## Acknowledgments

The authors acknowledge Haico van Attikum for providing U2OS GFP-RPB1 and U2OS osTIR1 cells, Leon Mullenders for providing CS1AN-SV40 cells expressing GFP or GFP-CSB, Rick Wood for his generous gift of XPA antibody, and Brian Magnuson valuable help with the BrU-seq analysis. This work was funded by an LUMC Research Fellowship and an NWO-VIDI grant (ALW.016.161.320) to MSL, a Leiden University Fund (LUF) grant to DvdH (W18355-2-EM), an NWO-Veni grant to CGS, an ERC grant to ACOV (310913), and a Dutch Cancer Society (KWF-Young Investigator Grant: 11367) to RG-P. The MV and JAM labs are part of the Oncode Institute, which is partly funded by the Dutch Cancer Society (KWF).

---

**Supplemental Figure 1.**
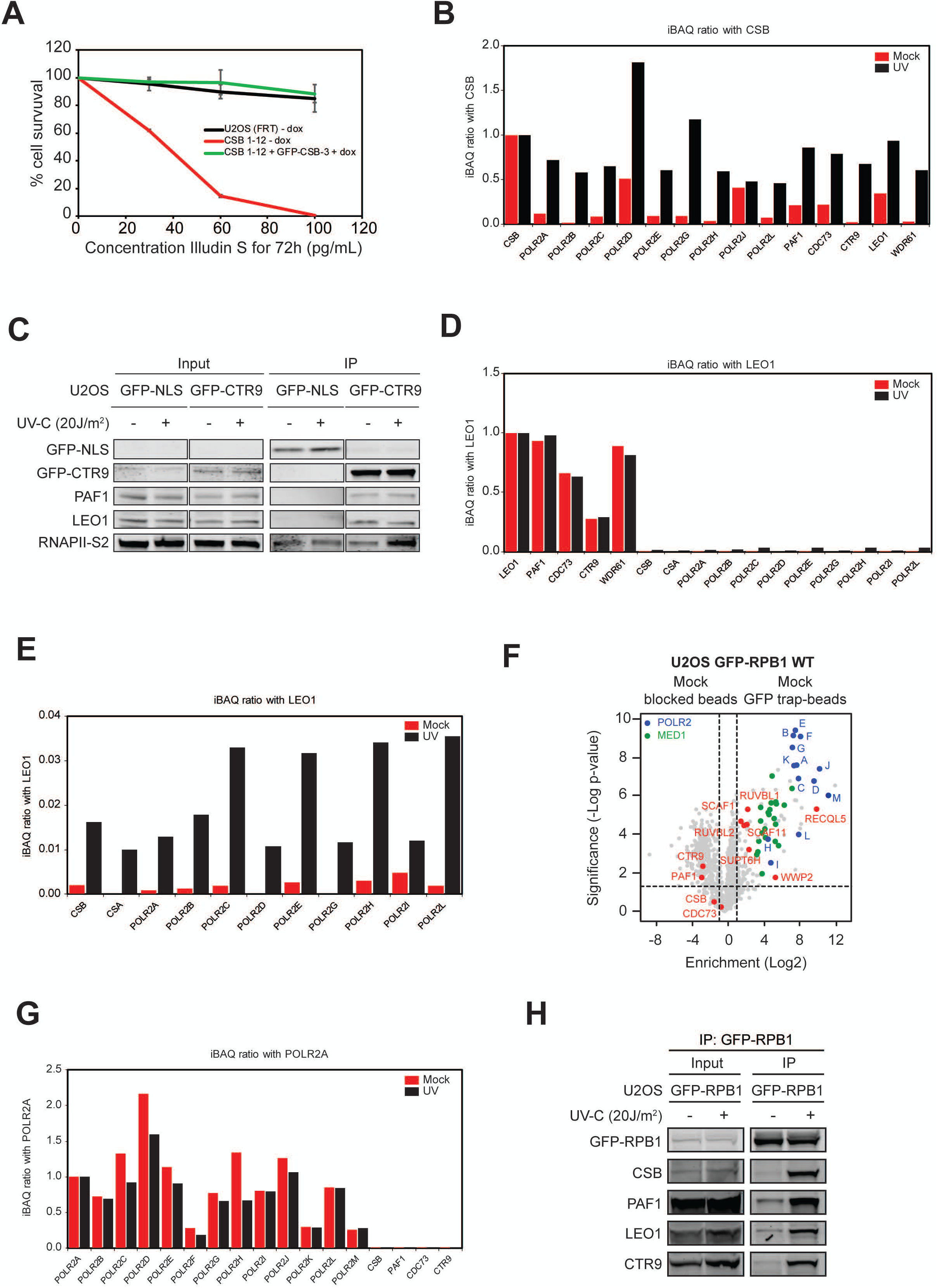
Interactions between CSB – RNAPII – PAF1C. (**A**) Clonogenic Illudin S survival of WT, CSB-KO and reconstituted CSB-KO with GFP-CSB. Data represent mean ± SEM of two independent experiments. (**B**) Stoichiometry of the interactions of the indicated proteins with GFP-CSB in unirradiated (red) and UV-irradiated U2OS CSB-KO cells (black) based on the iBAQ values obtained from label-free MS. (**C**) Co-immunoprecipitation of GFP-NLS or GFP-CTR9 from U2OS cells in the absence or presence of UV-induced DNA damage. (**D**) Stoichiometry of the interactions of the indicated proteins with GFP-LEO1 in unirradiated (red) and UV-irradiated RPE1-hTERT cells (black) based on the iBAQ values obtained from label-free MS. (**E**) Stoichiometry of the interactions of the indicated proteins with GFP-LEO1 in unirradiated (mock; red) and UV-irradiated U2OS cells (black) based on the iBAQ values obtained from label-free MS. (**F**) Volcano plot depicting the enrichment of proteins after pull-down of GFP-RPB1 from U2OS cells analysed by label-free MS. The enrichment (log^2^) is plotted on the x-axis and the significance (t-test −log^10^ p-value) is plotted on the y-axis. Highlighted are significantly enriched subunits of RNAPII (blue), mediator (green) and several known interactors of RNAPII (red). Note that CS proteins and PAF1C subunits are not detected in unirradiated cells under these conditions. (**G**) Stoichiometry of the interactions of the indicated proteins with POLR2A / RPB1 in unirradiated (red) and UV-irradiated cells (black) based on the iBAQ values obtained from label-free MS. (**H**) Co-immunoprecipitation of GFP-RPB1 from U2OS cells in the absence or presence of UV-induced DNA damage.

**Supplemental Figure 2.**
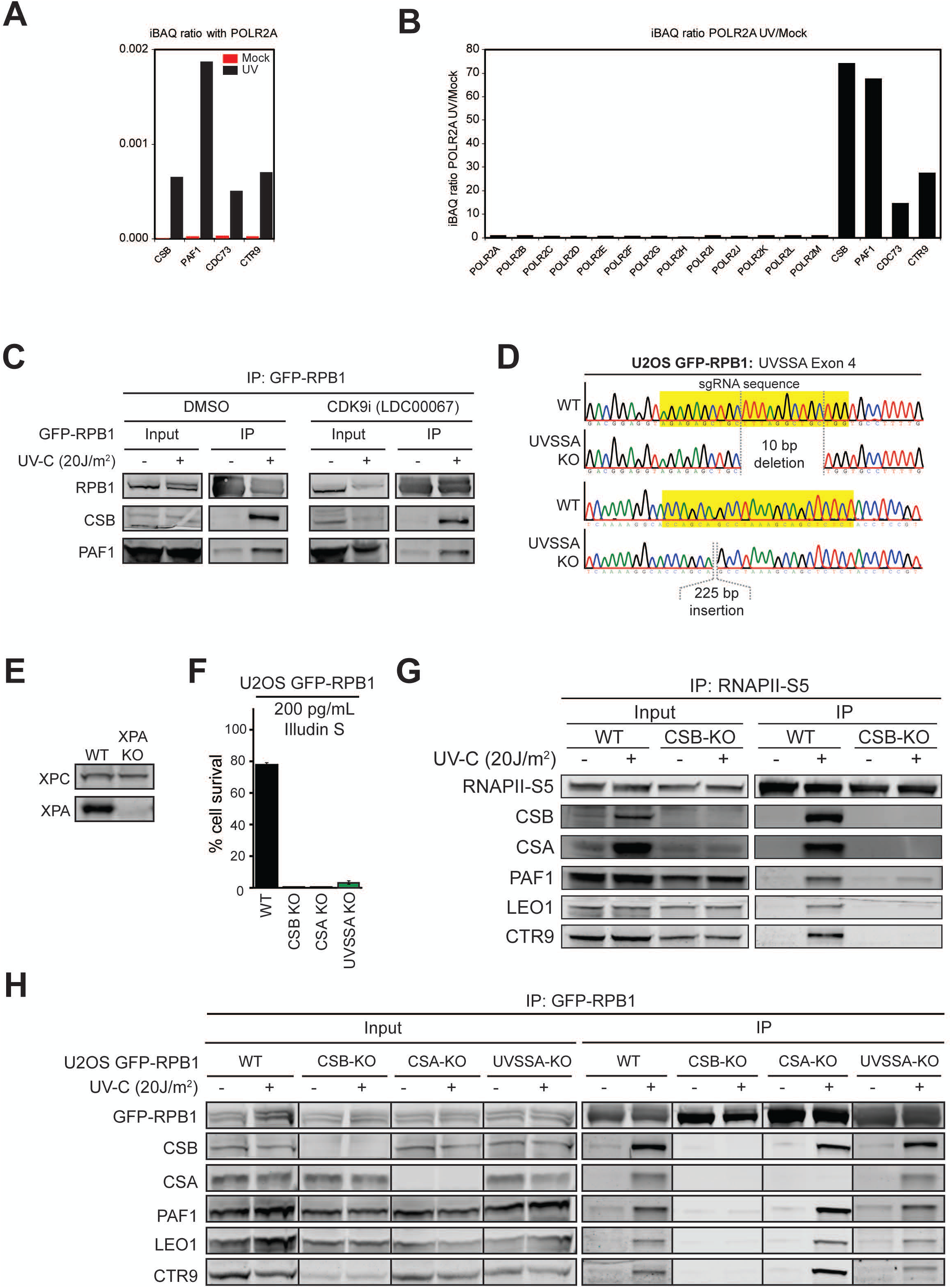
CSB mediates the UV-induced RNAPII – PAF1C interaction. (**A**) Stoichiometry of the interactions of CSB and PAF1C proteins with POLR2A / RPB1 in unirradiated (red) and UV-irradiated cells (black) based on the iBAQ values obtained from label-free MS. (**B**) The fold-increase of the CSB and PAF1C interaction with POLR2A / RPB1 after UV irradiation based on the iBAQ values. (**C**) Co-immunoprecipitation of GFP-RPB1 from U2OS cells treated with DMSO or with 10 µM CDK9 inhibitor (LDC00067) for 8 hrs in the absence or presence of UV-induced DNA damage. (**D**) Sequencing to confirm the knockout of *UVSSA* in U2OS GFP-RPB1 cells. (**E**) Western blot confirmed the knockout of XPA in U2OS cells. (**F**) Clonogenic Illudin S survival of GFP-RPB1 WT, CSA, CSB, and UVSSA knockout cell lines at a single dose. Data represent mean ± SEM of two independent experiments. (**G**) Co-immunoprecipitation of endogenous RNAPII using Ser5-specific antibodies from U2OS WT or CSB-KO cells in the absence or presence of UV-induced DNA damage. (**H**) Co-immunoprecipitation of GFP-RPB1 from U2OS WT, CSA, CSB, and UVSSA knockout cells in the absence or presence of UV-induced DNA damage.

**Supplemental Figure 3.**
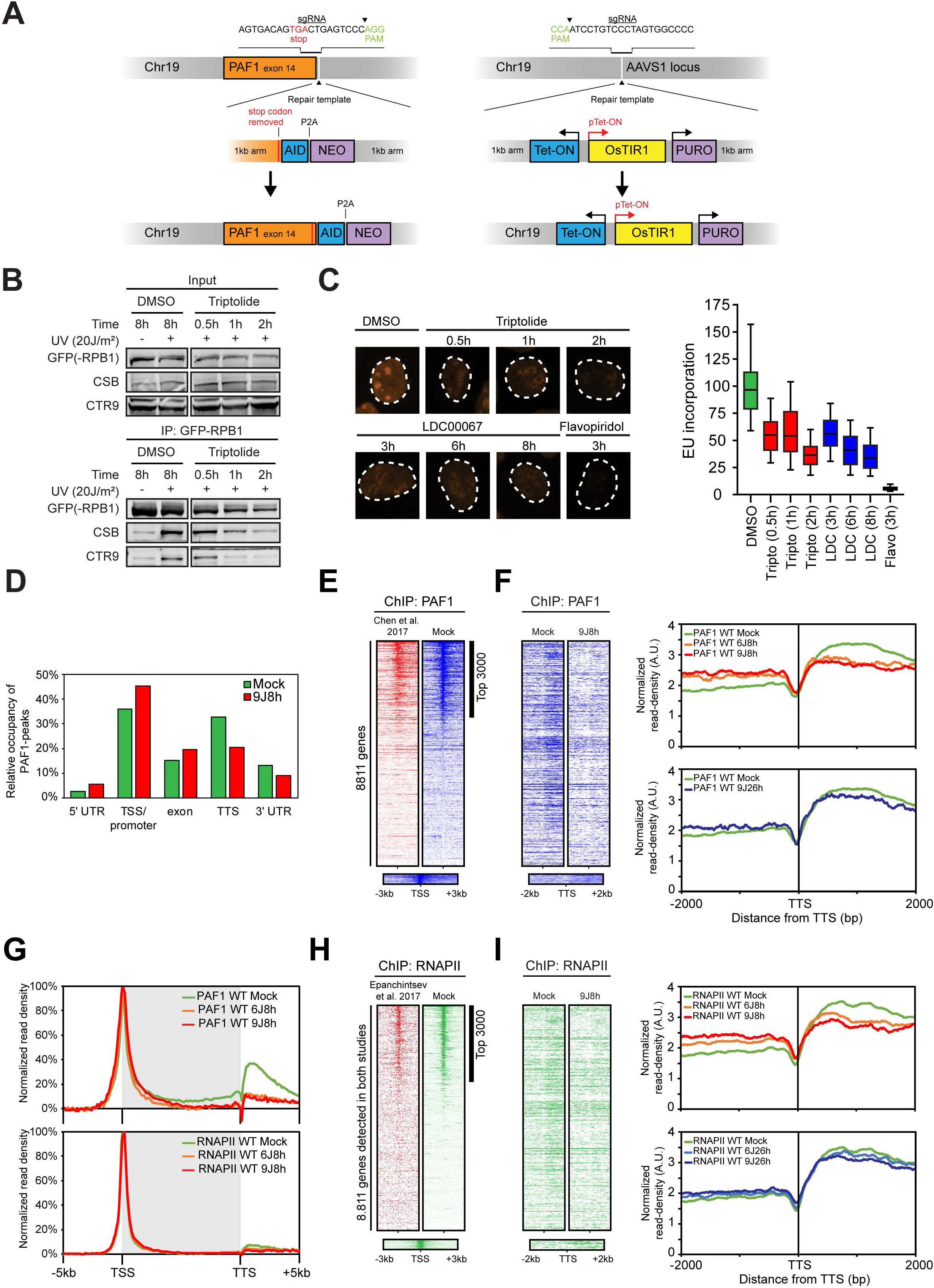
PAF1-AID knockin cells, transcription inhibitor controls, and comparison of ChIP-seq data with published datasets. (**A**) Outline of the approach to generate PAF1-AID knockin cells. The DOX-inducible osTIR1 cassette was first targeted to the *AAVS1* locus. The AID-NEO cassette was subsequently targeted to the last exon of the endogenous *PAF1* locus whereby the stop codon was removed to generate a PAF1-AID in-frame fusion. (**B**) Co-immunoprecipitation of GFP-RPB1 in the presence of 0.55 µM Triptolide (TFIIH/XPB inhibitor) for the indicated time, in unirradiated cells or 1h after UV irradiation during the incubation with inhibitors. (**C**) Representative images of U2OS cells treated with the indicated transcription inhibitors after pulse-labelling with 5-ethynyl-uridine (5-EU) and detection by Click-It chemistry. The right panel shows the quantification of the 5-EU signal. Data is represented in boxplots showing the median, 50% and 95% percentile of two independent experiments. (**D**) Occupancy of PAF1 binding-peakds across the coding regions of the genome in unirradiated (mock) and UV-irradiated cells. Peaks were identified using MACS2 of sample over input. (**E**) Heatmaps around the TSS from PAF1 ChIP-seq data of 8,811 genes ranked according to PAF1 signal at the TSS in unirradiated cells (mock; blue) of which the first 3,000 genes show strong PAF1 enrichment. Data is compared to publish data (in red) (**F**) Heatmaps and metaplots from PAF1 ChIP-seq data around the TTS of the 3,000 genes with strong PAF1 binding (identified in E and Figure 5F) in unirradiated (mock) or UV-irradiated cells (8h after 6J/m^2^ and 8 or 26h after 9J/m^2^). (**G**) Metaprofile of PAF1 (top) or RNAPII (bottom) binding across the top 3,000 genes with PAF1 binding at the TSS as identified in E and Figure 5F. (**H**) Heatmaps on genes as in E, but for RNAPII ChIP-seq data in unirradiated cells (mock; blue), compared to publish data (in red). (**I**) Heatmaps and metaplots on genes as in F, but for RNAPII ChIP-seq data in unirradiated (mock) or UV-irradiated cells (8h after 6J/m^2^ and 8 or 26h after 9J/m^2^).

**Supplemental Figure 4.**
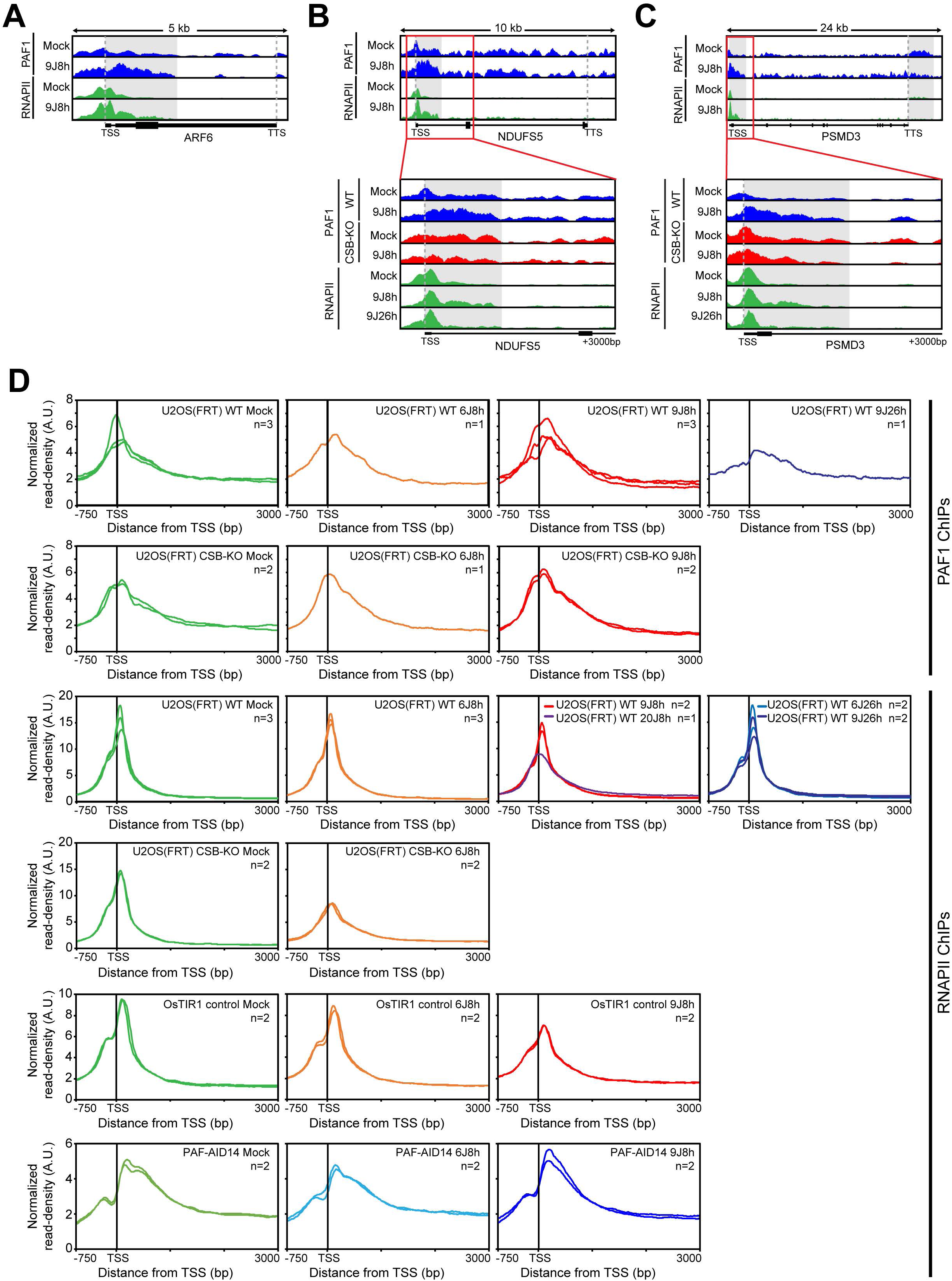
Distribution of PAF1 and RNAPPI across several representative genes and all individual ChIP-seq replicates. (**A-C**) UCSC genome browser track showing the PAF1 and RNAPII read densities across the (**A**) *ARF6* gene of 5 kb, (**B**) *NDUFS5* gene of 10 kb, and (**C**) *PSMD3* gene of 24 kb in U2OS WT cells. Insets of the first 3 kb of each gene is shown at the bottom for U2OS WT and CSB-KO cells. (**D**) Metaplots of all the individual PAF1 ChIP-seq and RNAPII ChIP-seq replicates of the top 3,000 genes around the TSS in unirradiated and UV-irradiated U2OS WT, U2OS CSB-KO, U2OS osTIR1, and U2OS PAF1-AID14 cells.

**Supplemental Figure 5.**
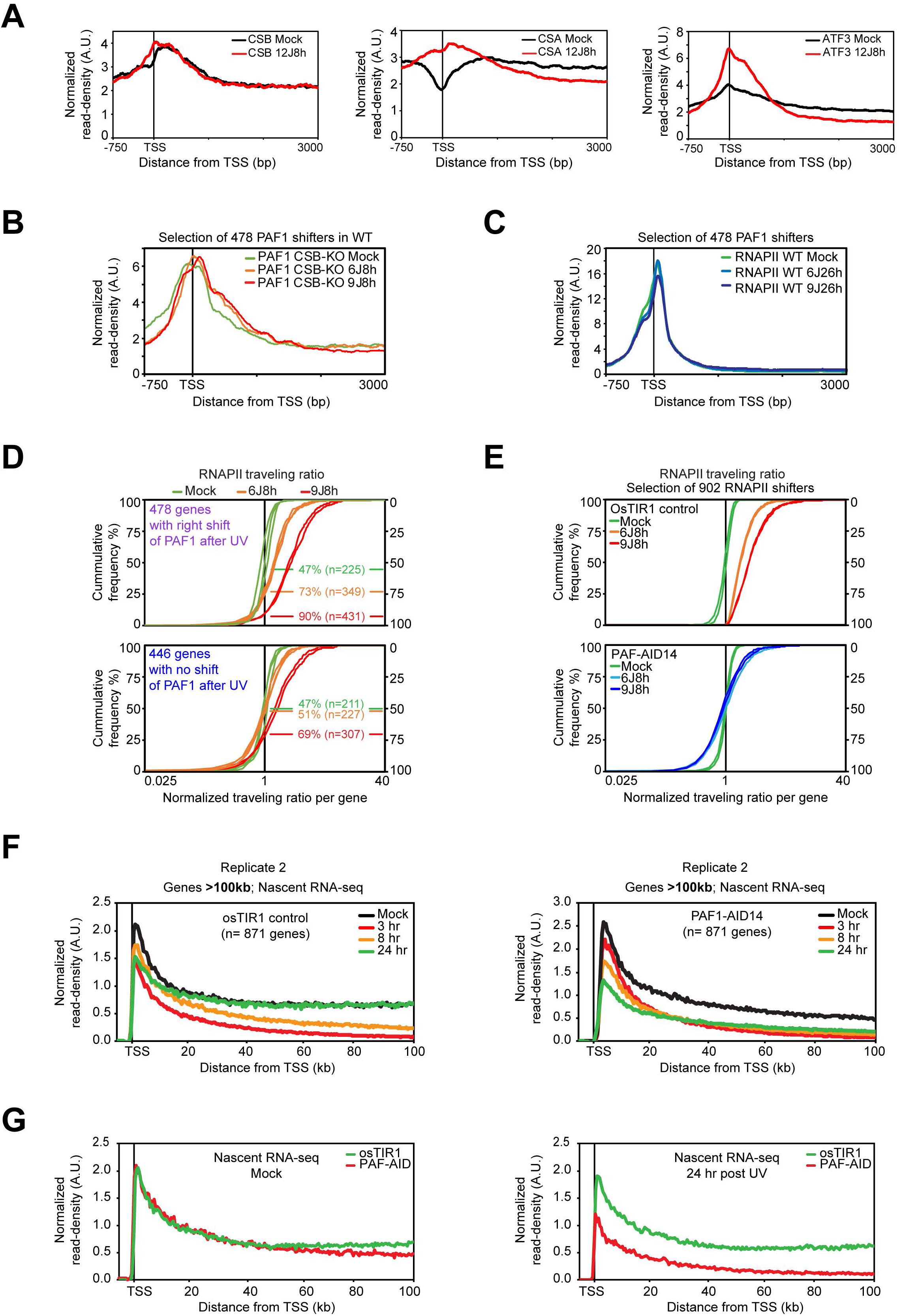
Metaplots around TSS and/or normalized traveling ratios of CSB, CSA, ATF3, PAF1 and RNAPII ChIP-seq and replicate validation of BrU-seq data. (**A**) Metaplots of CSB, CSA and ATF3 ChIP-seq from the Egly lab of the top 3,000 genes around the TSS in unirradiated and UV-irradiated cells (8 hrs after 12 J/m^2^). (**B**) Metaplots of PAF1 ChIP-seq in unirradiated and UV-irradiated CSB-KO cells (8 hrs after 6 J/m^2^ or 9J/m^2^) of the top 478 genes that show a strong UV-associated shift in PAF1 around the TSS in WT cells (Figure 6G). (**C**) Metaplots of RNAPII ChIP-seq in unirradiated and UV-irradiated WT cells (26 hrs after 6 J/m^2^ or 9J/m^2^) of the top 478 genes that show a strong shift in PAF1 around the TSS in WT cells (Figure 6G). (**D**) The ratio of the RNAPII traveling ratio for the PAF1 right-shift 478 genes (upper panel) or the no-shift 446 genes (lower panel) from Figure 6G, H relative to the average traveling ratio in the unirradiated control (set to 1). Shown are 3 independent replicates in unirradiated cells (green), and two replicates 8h after UV irradiation with 6 J/m^2^ (in orange) and two replicates 8h after 9 J/m^2^ (in red). The y axes indicate percent of all genes. Percentages and n indicated in the plot refer to the percentage and number of the 478 or 446 genes with a normalized traveling ratio above 1. (**E**) The ratio of the RNAPII traveling ratio for the 902 genes that show the strongest RNAPII shift in osTIR1 cells relative to the average traveling ratio in the unirradiated control (set to 1). The lower panel shows the ratio of the RNAPII traveling ratio of the same genes in PAF1-AID clone 14. Shown are 2 independent replicates in unirradiated cells (mock), and two replicates 8h after UV irradiation with 6 J/m^2^, and two replicates 8h after 9 J/m^2^. The y-axes indicate percent of all genes. (**F**) Independent replicate of the BrU-seq experiment shown in Figure 7B. Metaplots of nascent transcription in genes of >100 kb in either osTIR1 cells (left panel) or PAF1-AID cells (right panel) that were either mock-treated, or UV-irradiated (7 J/m^2^) and analysed at the indicated time-points (3, 8, or 24 hrs). The relative distribution of nascent transcript read density (in reads per thousand base-pairs per million reads) was normalized to the absolute nascent transcript intensities measured by RRS experiments that were performed in parallel to the BrU-seq experiments using the same cells and time-points (see Figure 4G, H and Figure 7B). (**G**) Metaplots of BrU-seq reads densities in unirradiated (left panel) or UV-irradiated (right panel, 24 hrs after UV) U2OS osTIR1 compared to PAF-AID14 cells for comparison from replicate 1 shown in Figure 7B.

**Supplemental Figure 6.**
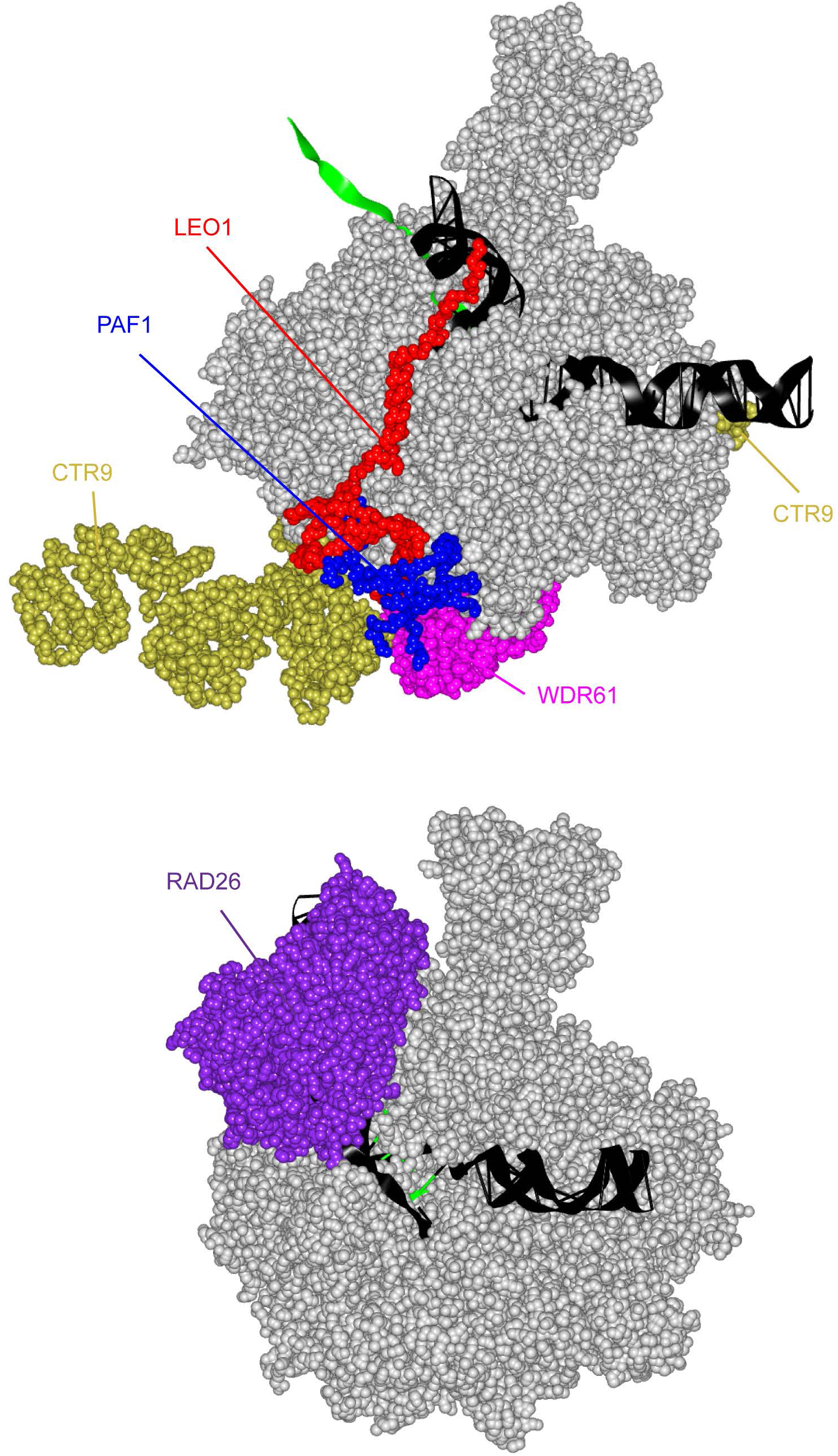
Structures of RNAPII with bound RAD26 or PAF1C complex. The upper panel shows the Cryo EM structure of RNAPII (in silver) bound to PAF1C subunits (color-coded as indicated in the figure) from the Cramer lab (PDB-ID: 6GMH). The lower panel shows the Cryo EM structure of RNAPII (in silver) bound to RAD26, which is the yeast orthologue of CSB, from the Wang lab (PDB-ID: 5VVR).

